# Structural dynamics of the intrinsically disordered linker region of cardiac troponin T

**DOI:** 10.1101/2024.05.30.596451

**Authors:** Jasmine Cubuk, Lina Greenberg, Akiva E. Greenberg, Ryan J. Emenecker, Melissa D. Stuchell-Brereton, Alex S. Holehouse, Andrea Soranno, Michael J. Greenberg

## Abstract

The cardiac troponin complex, composed of troponins I, T, and C, plays a central role in regulating the calcium-dependent interactions between myosin and the thin filament. Mutations in troponin can cause cardiomyopathies; however, it is still a major challenge to connect how changes in sequence affect troponin’s function. Recent high-resolution structures of the thin filament revealed critical insights into the structure-function relationship of troponin, but there remain large, unresolved segments of troponin, including the troponin-T linker region that is a hotspot for cardiomyopathy mutations. This linker region is predicted to be intrinsically disordered, with behaviors that are not well described by traditional structural approaches; however, this proposal has not been experimentally verified. Here, we used a combination of single-molecule Förster resonance energy transfer (FRET), molecular dynamics simulations, and functional reconstitution assays to investigate the troponin-T linker region. We show that in the context of both isolated troponin and the fully regulated troponin complex, the linker behaves as a dynamic, intrinsically disordered region. This region undergoes polyampholyte expansion in the presence of high salt and distinct conformational changes during the assembly of the troponin complex. We also examine the ΔE160 hypertrophic cardiomyopathy mutation in the linker and demonstrate that it does not affect the conformational dynamics of the linker, rather it allosterically affects interactions with other troponin complex subunits, leading to increased molecular contractility. Taken together, our data clearly demonstrate the importance of disorder within the troponin-T linker and provide new insights into the molecular mechanisms driving the pathogenesis of cardiomyopathies.

**SIGNIFICANCE STATEMENT:** Troponin plays a central role in regulating heart contraction, and mutations in troponin can cause human cardiomyopathies. There are several functionally-significant regions of troponin that have not been structurally resolved, including the troponin T linker region that contains multiple cardiomyopathy mutations. In these unresolved regions, it is not possible to understand how changes in sequence affect function. We used computational and experimental techniques to demonstrate that this linker is dynamic and intrinsically disordered both in isolation and in the fully regulated thin filament. Moreover, we show how a cardiomyopathy mutation in this region affects function via allosteric disruption of intermolecular interactions. Our results highlight the need to consider how key mutations affect troponin disorder rather than the structure-function relationship.

## INTRODUCTION

Familial cardiomyopathies, leading causes of heart failure and indicators for heart transplantation, are characterized by altered cardiac contractility and ventricular remodeling (1). Excellent clinical studies have defined the genetic landscape of cardiomyopathies, and they have shown that these diseases can be caused by point mutations in sarcomeric proteins involved in cardiac contraction, including cardiac troponin T (TnT) (2). While the association of troponin mutations with cardiomyopathies is unambiguous (3), hundreds of variants have been found in troponin, and it remains a challenge to determine whether or not a given patient-specific mutation is pathogenic. This has led to the classification of many variants of unknown significance (VUS), and as such, genetic information is not typically used to inform patient care (4).

At the molecular scale, cardiac contraction is powered by myosin motors in the sarcomere that pull on actin in regulated thin filaments (RTFs). Excellent biochemical and biophysical experiments have established that the interactions between myosin and actin are gated by calcium via the thin filament regulatory components troponin (a complex composed of troponins I, T, and C) and tropomyosin. During diastole (the cardiac cycle phase where the heart relaxes), intracellular calcium levels are low, and tropomyosin blocks the myosin strong-binding sites on actin (5–7). When intracellular calcium levels rise, calcium binds to troponin C, leading to alterations in the conformation of the troponin complex and repositioning of tropomyosin that enables the strong binding of myosin to actin. Recently, cryogenic electron microscopy (cryo-EM) enabled the visualization of the regulated thin filament (7–11). While some regions of troponin are clearly resolved, such as the troponin core and the tropomyosin overlap region of troponin T, there are large, functionally significant stretches of troponin that have not been resolved in any cryo-EM structure.

One critical unresolved region of the thin filament is the troponin-T linker region that connects the troponin core (where calcium binds) to tropomyosin. This linker is a hot spot for both VUS and pathogenic cardiomyopathy mutations (12–15). The linker is highly enriched in charged residues, and computational tools predict that it is likely an intrinsically disordered region (IDR) (16, 17); however, this prediction has not been tested experimentally, and recent work has called into question whether highly-charged blocky sequences such as the troponin linker are truly disordered (18). Compared to structured regions, IDRs tend to be more dynamic, exploring a large conformational space, and are thus not well described by traditional structural techniques (19, 20). Moreover, while computational tools have been developed to predict the potential pathogenicity of VUS in the clinic, these tools are generally based on the predicted effects of mutations on structured regions, making them poorly poised for classifying VUS in predicted IDRs, such as the troponin-T linker (21–24). There is thus an outstanding need to understand the structure and function of this linker region.

Here, we use a combination of computational and experimental approaches to investigate the linker region of troponin T both in isolation and in the context of the fully regulated thin filament. We use motility assays to test the effect of mutations on the mesoscopic scale. To capture the conformational changes and dynamics at the molecular level, we harnessed single-molecule fluorescence techniques, which have been used to provide valuable insights into the C-terminal region of troponin I (25, 26). Moreover, we apply these tools to investigate the effects of a pathogenic mutation within the troponin-T linker region, deletion of E160 (ΔE160).

## METHODS

### Protein expression, purification, and labeling

Cardiac actin and myosin were purified from cryoground porcine ventricles as previously described (27–30). Human cardiac tropomyosin was expressed recombinantly in E. coli and purified as previously described (27–30). For the ATPase and in vitro motility assays, human cardiac troponins I, T, and C were expressed recombinantly, complexed, and purified as previously described (31–33). For all single-molecule experiments, troponins I, T, and C were cloned into the pNH-TrxT vector (pNH-TrxT was a gift from Opher Gileadi (Addgene plasmid # 26106; http://n2t.net/addgene:26106; RRID:Addgene_26106, (34)) that contained an N-terminal thioredoxin, hexahistidine, and TEV cleavage site followed by troponin T. Troponin T has no native cysteines, so for the FRET experiments, cysteines for labeling were introduced into positions A153C and A213C flanking the troponin-T linker region using QuikChange mutagenesis (Agilent), and we will refer to this construct as **TnT_153,213_**. The ΔE160 mutant troponin T construct was also generated using QuikChange mutagenesis. Troponin subunits were then expressed in BL21(DE3)CodonPlus E. coli (Agilent) with IPTG induction and overnight expression at 18°C.

All purification steps were done at 4°C unless otherwise noted. To purify troponin subunits, cells were lysed with sonication in a buffer containing 20 mM MOPS pH 7, 20 mM Imidazole pH 7, 500 mM NaCl, 0.5 mM EGTA, 0.5% Igepal, 1 mM β-mercaptoethanol, 1 mM PMSF, 0.01 mg/ml aprotinin, 0.01 mg/mL leupeptin, and 6 M urea and then centrifuged at 164,000 g for 1 hour to separate the membrane and cytosolic fractions. The supernatant was purified by affinity chromatography using a His-Trap HP FPLC column (Cytiva) with Buffer A (same as lysis buffer minus Igepal) and eluted with 0-100% Buffer B (same as Buffer A plus 250 mM Imidazole). Fractions containing troponin were pooled and dialyzed overnight into cutting buffer containing 10 mM Imidazole (pH 7), 20 mM MOPS (pH 7), 250 mM NaCl, 1 mM DTT, 0.5 mM EGTA, and 2 M urea. The concentration of hexahistidine-thioredoxin-TEV-troponin was measured spectroscopically. The hexahistidine-thioredoxin-TEV-troponin was cleaved using a His-TEV enzyme at a 1:20 mass ratio to troponin. The His-TEV enzyme and troponin were incubated at 37°C for 2 hours, then centrifuged for 10 minutes at 4,000 g, and the supernatant was clarified through a 0.45 µm syringe filter. The supernatant was then loaded onto a His-Trap HP FPLC column, and the flow through (which did not bind to the column) was collected, concentrated using Amicon Ultra Centrifugal Filters (Millipore), flash frozen in liquid nitrogen, and stored at minus 80°C. All protein concentrations were determined spectroscopically using the molecular weight to predict extinction coefficients in https://web.expasy.org/protparam/.

His-tagged TEV protease (pDZ2087, a gift from David Waugh (Addgene plasmid # 92414; http://n2t.net/addgene:92414; RRID:Addgene_92414)) was expressed in BL21(DE3)CodonPlus E. coli with IPTG induction and overnight expression at 18°C. His-TEV was purified by following the method described in (35) with modification. Briefly, bacterial cell pellets were lysed by sonication in 20 mM HEPES pH 7.5, 10 mM Imidazole, 300 mM NaCl, 0.5% Igepal, 1 mM β-mercaptoethanol,1 mM and PMSF. The protein was purified using a His-Trap HP FPLC column (Cytiva) followed by gel-filtration chromatography over a Superdex 16/600 HL FPLC column (Cytiva) in buffer containing 20 mM HEPES, 250 mM NaCl, 2 mM EDTA, 1 mM DTT. The fractions containing His-TEV were pooled and dialyzed overnight into buffer containing 20 mM HEPES, 20 mM NaCl, 2 mM EDTA, and 1 mM DTT. The protein was concentrated to 2.5 mg/mL, flash frozen in liquid nitrogen, and stored at minus 80°C.

**TnT_153,213_** was labeled with Alexa Fluor 488 maleimide (Molecular Probes) under denaturing conditions in buffer A (50 mM HEPES pH 7.3, 6 M Urea) at a dye/protein molar ratio of 0.7/1 for 16 hours at 4°C. Single-labeled protein was isolated via ion-exchange chromatography (Mono Q 5/50 GL, GE Healthcare - protein bound in buffer A (+5 mM BME) and eluted with 10-40% buffer B (buffer A + 1 M NaCl) gradient over 60 min) and UV-Vis spectroscopic analysis to identify fractions with 1:1 dye:protein labeling. Single-labeled Alexa Fluor 488 maleimide labeled troponin was then subsequently labeled with Alexa Fluor 594 maleimide at a dye/protein molar ratio of 1.5/1 for 2.5 hours at room temperature. Double-labeled (488:594) protein was then further purified via ion-exchange chromatography (Mono Q 5/50 GL, GE Healthcare).

### Single-molecule fluorescence

Single-molecule FRET measurements were performed on both a modified Picoquant MT200 instrument (Picoquant, Berlin, Germany) and an equivalent custom-built instrument. Specifications of the instrumentation were previously published (36), and we briefly summarize them here. Pulsed Interleaved Excitation (PIE) is generated by alternating emission from a diode laser (LDH-D-C-485, PicoQuant, Germany) and a supercontinuum laser (SuperK Extreme, NKT Photonics, Denmark) filtered by a z582/15 band pass filter (Chroma) with an excitation rate of 20 MHz and a delay between the two pulses of ∼ 25 ns. Laser beams were focused through a 60x 1.2 N.A. UPlanSApo Superapochromat water immersion objective (Olympus, Japan), which also collects emission. Emitted photons were filtered through a dichroic mirror (ZT568rpc, Chroma, USA), a long pass filter (HQ500LP, Chroma Technology), and the confocal pinhole (100 μm diameter) and, subsequently, selected according to polarization. Donor and acceptor photons were first separated by a dichroic mirror (585DCXR, Chroma) and further refined by using band pass filters, ET525/50m or HQ642/80m (Chroma Technology), respectively. Finally, photons were focused on SPAD detectors (Excelitas, USA), and arrival times were recorded using a HydraHarp 400 TCSPC module (PicoQuant, Germany). Experiments were conducted at 70 μW of donor excitation as measured at the back aperture of the objective, and the acceptor excitation was adjusted to match the total emission intensity after donor excitation. Experiments were conducted at a protein concentration of 100 pM (estimated from dilutions of samples with known concentration based on absorbance measurements) in 50 mM HEPES pH 7.4, 200 mM β-mercaptoethanol, and 0.001% Tween 20, at a room temperature of 295 ± 0.5 K. Temperature experiments were performed using 50 mM HEPES pH 7.4, 25 mM KCl, 4 mM MgCl_2_ and 2 mM EGTA. Binding experiments were performed with either 50 mM HEPES pH 7.4, 25 mM KCl, 4 mM MgCl_2_, 2 mM EGTA (absence of CaCl_2_, 99.96% chelated to EGTA even if 1 mM residual CaCl_2_ were present in the solution according to MaxChelator (37)) or 50 mM HEPES pH 7.4, 25 mM KCl, 4 mM MgCl_2_, and 2 mM CaCl_2_ (which we will refer to as “+2 mM CaCl_2_” condition). The second condition was chosen to match the *in vitro* motility assay conditions. Other additional components (e.g., salt and denaturant) are specified in the text when used.

Single-molecule FRET data were analyzed using the “Fretica’’ package for Mathematica developed by Daniel Nettels and Ben Schuler (University of Zurich, Zurich, CH). Data analysis was further processed in Mathematica (Princeton, NJ).

All single-molecule measurements are performed at least in duplicate (independent repeats) and reported errors are standard deviation from the multiple repeats, unless differently mentioned. Further details on the analysis are provided in **Supplementary Information**.

### Coarse-grained simulations

Coarse-grained molecular dynamics simulations of full-length cardiac troponin T were performed using the Mpipi-GG force field with the LAMMPS simulation package (38, 39). Mpipi-GG is a fine tuned version of the Mpipi (V1) force field, which was developed explicitly for the simulation of intrinsically disordered proteins and protein regions (40). All simulations were performed with 150 mM implicit salt concentration in the canonical (NVT) ensemble using a target temperature of 300 K. Simulations were minimized for 1,000 iterations or until the force tolerance was below 1 × 10^−8^ (kcal mol^−1^) per Å. The simulation temperature was maintained with a weakly-coupled Langevin thermostat with an integration timestep of 10 fs for all production runs. Simulations were performed with periodic boundary conditions in a 500 Å^3^ cubic box. Each independent simulation was run for 2 µs, with output coordinates saved every 0.1 ns.

All single-chain simulations were initially equilibrated for 5 ns and structures from this equilibration period were discarded. Prior structural characterization of troponin T has indicated two alpha helices (41). Based on helicity predictions, extant circular dichroism data, and prior biophysical characterization, we estimated helix 1 (residues 88-150) should be present while helix 2 (residues 228-267) should be present at about 30% population (41). With this in mind, we performed simulations to match this distribution. The final ensemble analyzed contained 85,504 conformers from six independent simulations reporting on an aggregate of ∼12 µs of simulation data. Simulations were analyzed using MDTraj and SOURSOP (42, 43).

Thin filament simulations were performed using the Mpipi force field with the LAMMPS simulation package (39, 40). All simulations were performed with 150 mM implicit salt concentration in the canonical (NVT) ensemble using a target temperature of 300 K. Simulations were minimized for 1,000 iterations or until the force tolerance was below 1 × 10^−8^ (kcal mol^−1^) per Å. The simulation temperature was maintained with a weakly-coupled Langevin thermostat with an integration timestep of 5 for all production runs. Simulations were performed with periodic boundary conditions in a 800 Å^3^ cubic box. Each independent simulation was run for 1 µs, with output coordinates saved every 0.05 ns. Construction of the thin filament for simulations was based on an cryoelectron microscopy structure (RCSB PDB structure 6KN7, (7)). The 6KN7 structure was coarse-grained using alpha carbons. All residues in the complex were fixed during simulations with the exception of the IDRs in TnT (P45379) and TnI (P19429). For the simulations, IDRs were inferred based on residues missing in the cryoelectron microscopy structure. For TnT, residues corresponding to likely IDRs and thus not fixed during simulation included residues 1-86, 151-198, and 273-288 in chains designated chain ‘T’ and chain ‘a’ in 6KN7. For TnI, residues corresponding to the IDR were residues 1-40 in chains designated as chain ‘U’ and chain ‘b’ in 6KN7. Initial starting configurations for the IDRs were constructed by iteratively placing beads using a random-walk algorithm. For terminal IDRs (all IDRs except for 151-198 in TnT), the random-walk started with placing a residue 3.81 Å from the closest possible position in the structure (for example, for IDR 1-86 in TnT, residue corresponding to residue 86 was placed 3.81 Å away from residue 87) and then iteratively placed residues at random locations 3.81 Å away from the previous residue with a requirement that and added residue could not be within a clashing distance of any existing residue. For IDR 151-198 in TnT, a similar strategy was followed except an additional constraint was added in that each additional residue after residue 151 had to be within the volume of a sphere with a size that was iteratively reduced such that the final residue added (residue 198) was approximately 3.81 Å from residue 199. Because IDRs were initially started in a ‘random walk’ configuration, simulations were allowed to equilibrate for 5 ns before data was collected to allow the IDRs to adopt appropriate configurations.

### In vitro motility assays

In vitro motility assays using regulated thin filaments were conducted as previously described (28, 33, 44). Briefly, flow chambers were assembled between two coverglasses. “Dead head” myosins were removed by ultracentrifugation in the presence of ATP and actin. Flow chambers were loaded as previously described. The final activation buffer contained 25 mM KCl, 60 mM MOPS pH 7.0, 2 mM EGTA, 4 mM MgCl_2_, 1% methyl cellulose, 1 mM DTT, 1 mg/mL glucose, 192 U/mL glucose oxidase, 48 μg/mL catalase, 1 μM troponin and tropomyosin, and varying calcium concentrations (pCa 4-9). The concentration of free calcium was balanced using MaxChelator (37). Videos were recorded at room temperature. Filaments were manually tracked using the ImageJ plugin MTrackJ (45). All data were collected from at least 3 separate days of experiments, and 25 filaments were tracked per condition. Statistical testing was done using a Mann-Whitney test.

### Regulated thin filament ATPase assays

The steady-state ATPase rate of myosin in the presence of actin, troponin, and tropomyosin was measured using an NADH-enzyme coupled assay as previously described (32, 46). Briefly, to polymerize G-actin to F-actin and to remove residual ATP, actin was dialyzed overnight into ATPase assay buffer containing 20 mM imidazole pH 7.5, 10 mM KCl, 2 mM MgCl_2_, and 1 mM DTT. The polymerized actin was stabilized by adding a 1.1x molar excess of phalloidin. Tropomyosin was combined with DTT and heated at 56°C for 5 minutes to reduce it and any aggregates were removed by centrifugation at 434,500 g for 30 minutes. The following components were combined in ATPase assay buffer containing 2 mM EGTA and 4 mM NTA: 1 µM S1 myosin, 3.5 µM phalloidin stabilized actin, 2 µM troponin, 1 µM tropomyosin, 0.313 mg/mL NADH (Sigma N7410-15VL), 0.5 mM Phospho(enol) Pyruvate (Sigma P0564), 1,000 U/mL Pyruvate Kinase (Sigma P9136), 200 U/mL Lactate Dehydrogenase (Sigma L1254), and the desired concentration of free calcium. The concentration of free calcium was calculated using MaxChelator (37). The reaction was initiated by the addition of 2 mM ATP. Experiments were carried out in a BioTek Synergy H1M plate reader in 96 well plates at 30°C with shaking. The absorbance at 340 nm was measured for 10 minutes at a continuous rate and the absorbance decreased linearly. The rate of change in the absorbance was measured over a range of calcium concentrations. Five technical repeats from at least 2 separate protein preparations were used. Data were fitted with a Hill equation, where the ATPase rate V is given by:

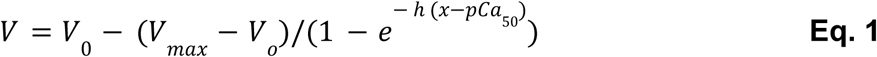

where *x* is the concentration of free calcium, *V*_0_ is the basal ATPase rate under fully inhibited conditions, *V*_*max*_ is the rate at fully activating conditions, ℎ is the Hill coefficient, and the *pCa*_50_ is the calcium concentration at which half-maximal activation is achieved. Uncertainties in the fitted parameters represent 95% confidence intervals, and statistical testing was performed using a Mann-Whitney test.

### Measurement of calcium binding to troponin C

Calcium binding to troponin C was measured spectroscopically as we have done previously (30, 47). Briefly, troponin C was mutated to remove native cysteines, and a cysteine was introduced into the calcium binding site (C35S, T53C, C84S). The protein was expressed in E. coli, purified, labeled with 2-(4′-(iodoacetamido)anilino)napthalene-6-sulfonic acid (IAANS; Toronto Research Chemicals) (48), and then reconstituted into troponin complexes as previously described (30, 47). Thin filaments consisting of 2 μM phalloidin stabilized actin, 0.45 μM tropomyosin, and 0.15 μM labeled troponin complex were reconstituted in a buffer (10 mM MOPS pH 7.0, 150 mM KCl, 3 mM MgCl2, 2 mM EGTA, and 1 mM DTT). A Fluoromax Spectrophotometer was used to excite the fluorophore at 330 nm, and emission from 370 to 550 nm was collected and integrated. Calcium chloride was added using a Microlab 600 Controller (Hamilton) and the concentration of free calcium was calculated using MaxChelator (37). The relative change in fluorescence was measured and fitted with a Hill Equation to extract the calcium concentration necessary for half saturation (i.e., pCa_50_). Data consisted of 5 technical replicates. Statistical testing was done using a Mann-Whitney test.

## RESULTS

The troponin-T linker, spanning from residues 150 to 203, connects the troponin core complex to tropomyosin (**Fig. 1**). The computational tool Metapredict (49, 50) predicts that this region may be intrinsically disordered. Consistent with this notion, recent cryo-EM high resolution structures of the thin filament do not report a high electron density for this region and no structural features have been resolved (7–11). Intrinsically disordered regions are indeed flexible, dynamic, and lack stable secondary structure, making them elusive to many structural biology methods. To test whether the troponin-T linker is indeed flexible and dynamic and how its sequence determines its conformational ensemble, we used single-molecule Förster Resonance Energy Transfer (smFRET). This approach enables quantification of the linker region conformations and dynamics in the contexts of both full-length troponin T protein and the full troponin complex. To this end, we expressed a cardiac troponin T full-length construct with cysteines flanking the linker region, in position 153 and 213, which we will refer to as **TnT_153,213_**. We then labeled these cysteine residues with the dye pair Alexa Fluor 488 and 594 (see **Methods**). The labeling efficiency was determined by Pulsed Interleaved Excitation (PIE) (51, 52), demonstrating that single-molecules of **TnT_153,213_**have been labeled with both one acceptor and one donor (**Fig. S1**).

**Figure 1:**
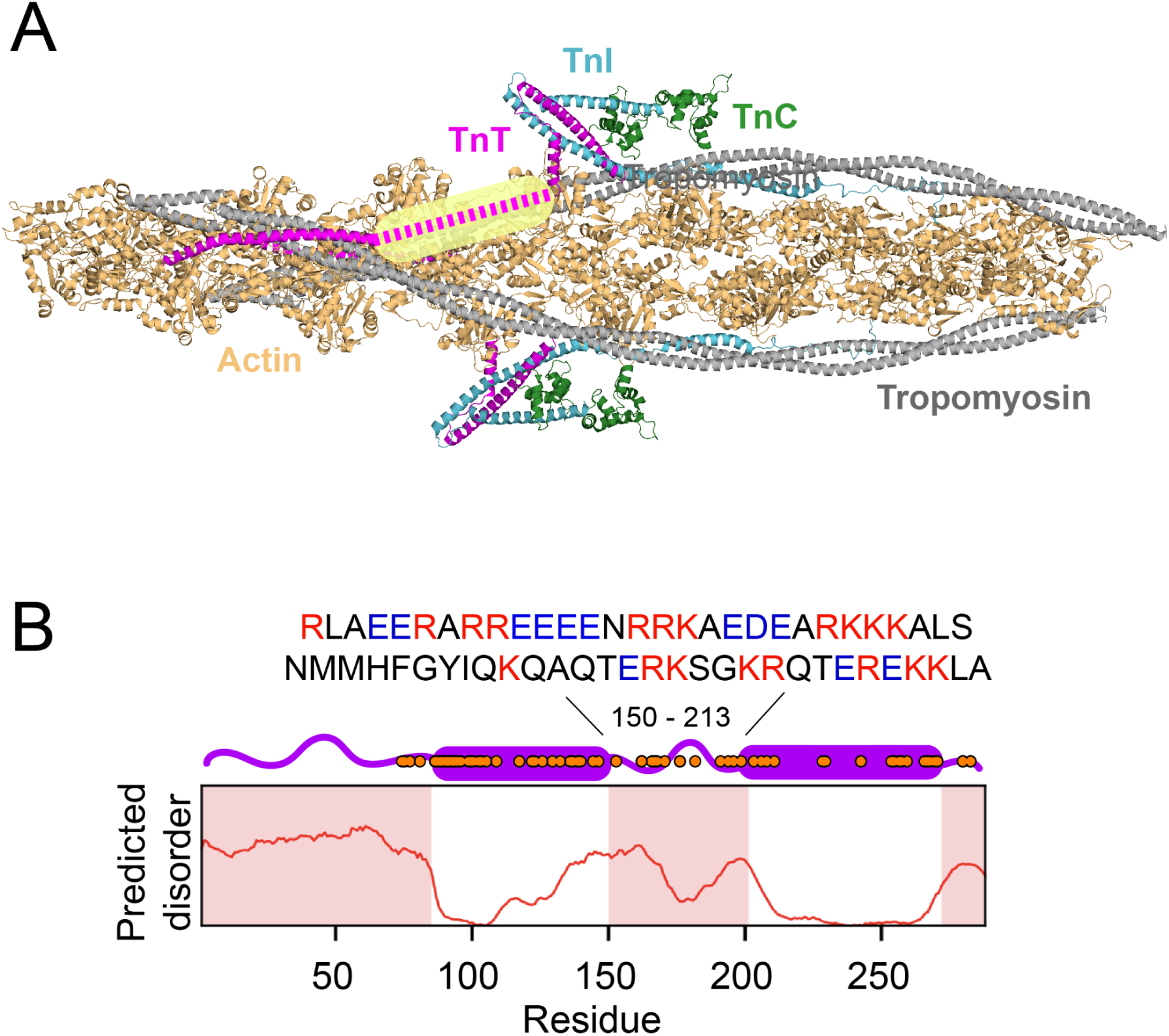
TnT sequence, linker. **A.** Representation of the thin-filament, with actin (light orange), tropomyosin (light gray), and the troponin complex with troponin C (TnC, green), troponin I (TnI, light blue), and troponin T (TnT, purple), based on PBD 6KN7. The dashed region in troponin T represents the unresolved linker region. **B.** Troponin T cartoon representation with the linker sequence highlighted (positive residues colored red, negative residues colored blue). Plot of predicted disorder shows the N terminal domain, linker, and C terminal domain to contain disorder. Orange dots represent pathogenic and likely pathogenic mutations found in troponin T, as identified via ClinVar.

### smFRET confirms that the troponin-T linker is intrinsically disordered

IDRs in solution are flexible and dynamic, exploring an ensemble of conformations, and they show monotonic changes in structure upon denaturation due to the lack of secondary structure. To determine whether troponin-T displays the hallmarks of an IDR, first, we performed confocal single-molecule FRET (smFRET) measurements of the freely-diffusing **TnT_153,213_** construct in isolation. The histogram of measured transfer efficiencies clearly demonstrates a single, well-defined peak associated with the donor-acceptor population (**Fig. 2 and Fig. S1)**. The measured distribution of transfer efficiencies (**Fig. 2A**) is close to the shot-noise limit (53, 54), indicating that the width of the distribution arises simply from the statistical uncertainty in the estimate of a single transfer efficiency rather than the distribution of distances sampled by the probes. The distribution shows a single characteristic mean transfer efficiency which implies that the linker region either: 1) adopts a single configuration corresponding to this transfer efficiency or 2) samples many configurations on a time scale much faster than the interphoton time such that the observed transfer efficiency is the average across the conformational ensemble.

**Figure 2:**
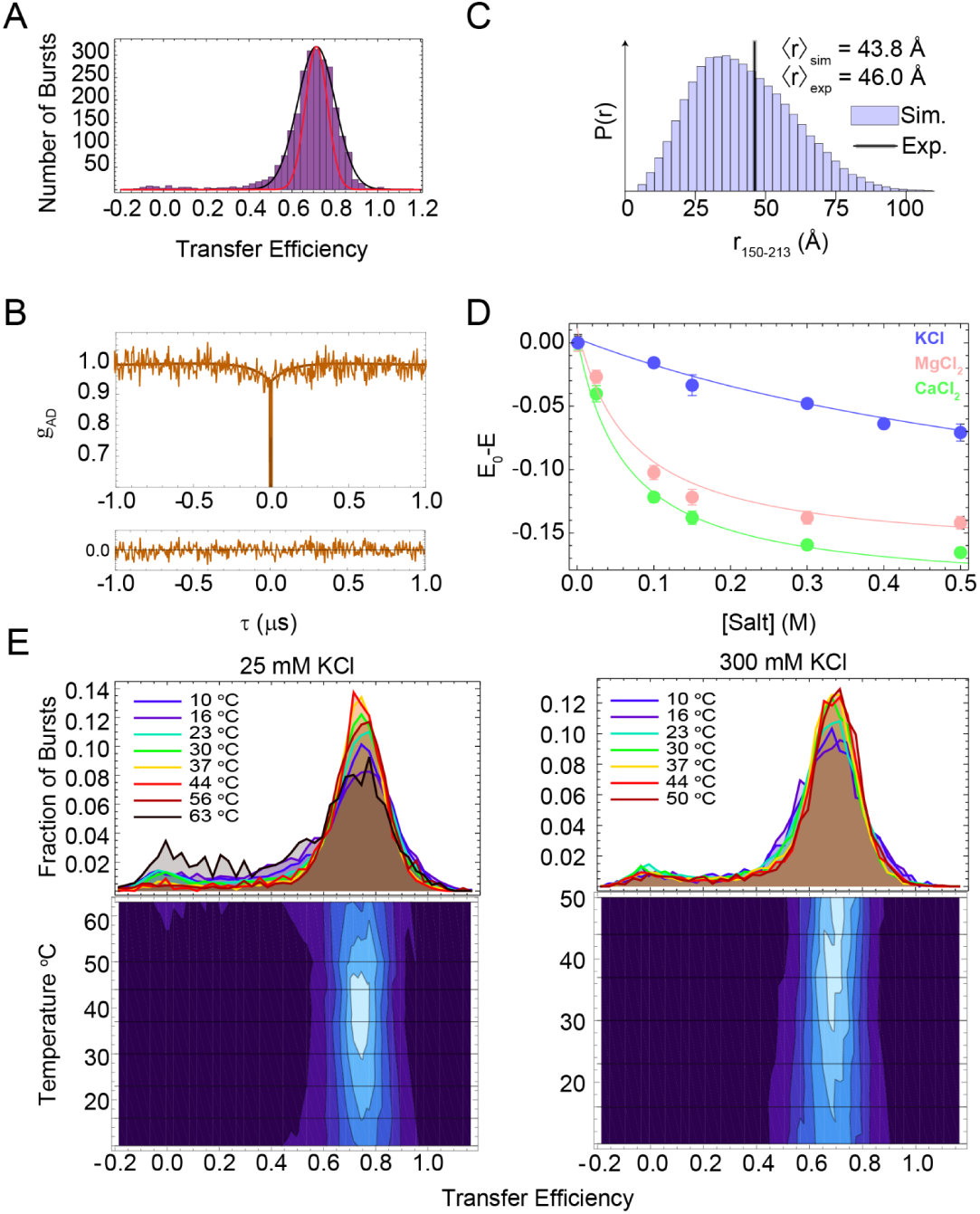
smFRET, nsFCS, and simulations of the troponin T-linker in isolation: **A.** Representative histogram of **TnT_153,213_** shows a single narrow distribution close to the shot-noise limit (red fit). We attribute the extra-broadening to the permutation of labeling positions. **B.** Nanosecond FCS (ns-FCS) acceptor-donor cross-correlation shows an anticorrelated amplitude, consistent with a dynamic linker. **C.** Coarse-grained simulations of full-length TnT were performed with the inter-residue distance between residues 153 and 213 reported here. Root-mean-squared distance from smFRET experiments is overlaid. **D.** Plot of change in transfer efficiencies as a function of salt (blue: KCl, pink: MgCl_2_, green: CaCl_2_). Values are mean and standard deviation of multiple (at least two) measurements. **E.** Top: Normalized histograms of **TnT_153,213_**FRET efficiency as a function of increasing temperature (10-63℃). Bottom: 2D plot of temperature and transfer efficiency in low salt (25 mM KCl, left) and high salt (300 mM KCl, right).

To discriminate between these two scenarios, we used nanosecond fluorescence correlation spectroscopy (nsFCS) to probe fast dynamics of the linker region (55, 56). nsFCS measures dynamics by quantifying the characteristic timescale associated with fluctuations in the distance sampled by the dyes. In the case of a dynamic linker, the cross-correlation of the donor-acceptor fluorescence would show an increase in amplitude over time due to the anticorrelated nature of donor and acceptor emission in the FRET process. Conversely, if the linker were not dynamic, the signal would show a flat correlation. Consistent with the notion that **TnT_153,213_** is dynamic, the cross-correlation function shows clear anticorrelation with a characteristic timescale of 120 ± 10 ns (**Fig. 2**). Taken together, our results demonstrate that the mean transfer efficiency seen for the troponin-T linker in solution corresponds to a fast reconfiguring dynamic ensemble, consistent with the notion that the linker is flexible and dynamic.

To test whether the linker contains any signature of stable secondary structure or whether it is affected by interactions with surrounding domains, we performed a chemical denaturation of **TnT_153,213_** with guanidinium chloride (GdmCl). A structured region would show discontinuous smFRET shifts due to local unfolding of the structure, whereas an IDR would show continuous shifts towards lower transfer efficiencies due to chain expansion. We observed a continuous shift toward lower transfer efficiencies with increasing concentrations of denaturant, which reflects an expansion of the linker and is a typical hallmark of a largely (if not completely) disordered sequence. Moreover, the monotonic trend of the expansion as a function of GdmCl in the absence of any discontinuities (**Fig. S2**) is consistent with undetectable contributions of secondary structure in the linker. These data further support that the troponin-T linker is likely an IDR.

Since the linker is sampling a dynamic distribution of distances, we can approximate the underlying distribution of distances using simple polymer models to compute the corresponding root-mean-square inter-residue distance. Using a Gaussian chain model, we obtained root-mean-square distance between the probes of 46 ± 0.5 Å (corrected for the dye linkers (57–59)).

To gain further details into the structure of the linker region, we performed coarse-grained molecular dynamics simulations of full-length troponin T with residual helicity encoded into the underlying simulation (see **Methods**). These simulations provided extensive sampling of the IDR conformational ensemble. We computed the distribution of distances sampled by positions 153 and 213 from the resulting ensemble (**Fig. 2C**). Note that the width of the simulated distribution of distances cannot be directly compared to the width of the distribution of single molecule transfer efficiencies, since the chain samples many configurations on a fast time scale and the experimentally measured mean transfer efficiency only reports the root-mean-square value of the underlying distribution. Therefore, we determined the root-mean-square inter-residue distance from the simulation, obtaining a value of 43.8 Å, which is in good agreement with our smFRET experiments. Taken together, our measurements, modeling, and simulations clearly demonstrate that the troponin-T linker is a dynamic, flexible, IDR in the context of the isolated full-length monomeric protein.

### Role of electrostatics and temperature on the linker conformation

Having established that the troponin-T linker is an IDR, we set out to determine the molecular driving forces underlying its disorder by examining the contributions of electrostatics and temperature in tuning this behavior. Given the concentration of both positively and negatively charged residues between the labeling positions, we expect that the linker would exhibit a typical polyampholyte behavior, where positive and negative charges within the chain contribute to attractive interactions and result in chain compaction (compared to an uncharged chain). In this scenario, we expect that the addition of salt would result in chain expansion due to electrostatic screening of charges within the chain. smFRET measurements of **TnT_153,213_**with increasing concentration of KCl resulted in a decrease in smFRET, corresponding to an expansion of the chain and confirming the role of electrostatics in modulating the conformational ensemble of the troponin-T linker (**Fig. 2**). Interestingly, while both monovalent (KCl) and divalent ions (MgCl_2_ and CaCl_2_) cause chain expansion, they do so to different extents. At 0.5 M salt, KCl shifts the root-mean-square inter-residue distance up to 4.92 ± 0.04 nm, while MgCl_2_ and CaCl_2_ shift it up to 5.43 ± 0.01 nm and 5.55 ± 0.01 nm, respectively.

Finally, we measured the temperature dependence of the linker dimensions. Based on the density of charged residues in the linker sequence and their significant change in the free energy of solvation with temperature, we expected to observe a further compaction of the chain as reported for other charged disordered sequences (60–62). Surprisingly, we did not observe any significant shift in the mean transfer efficiency between 10 and 63 °C, indicating that the linker is largely insensitive to temperature (**Fig. 2**).

### Linker conformations upon assembly of the troponin complex

Given that troponin T functions in the context of a complex consisting of troponins T, I, and C, we examined the contributions of these intermolecular interactions on the conformation of the linker. We purified human troponins I and C, mixed them with labeled **TnT_153,213_**, and measured the conformation of the linker region using smFRET. Given the nanomolar affinity that troponin I, T, and C have for one another, single-molecule spectroscopy provides an unique advantage in dissecting the gradual binding of troponin T.

First, we titrated an increasing concentration of unlabeled troponin I in the absence of troponin C (**Fig. S3**). We observed a clear shift in the mean transfer efficiency of **TnT_153,213_** toward lower values, which saturates above 10 nM of troponin I. The shift direction is consistent with an expansion of the linker when complexing with troponin I. The transfer efficiency change can be fitted to extract a binding affinity of troponin T for troponin I, which results in a K_D_ of 1.1 ± 0.2 nM in the absence of CaCl_2_ and a K_D_ of 0.7 ± 0.3 nM in the presence of 2 mM CaCl_2_.

When performing the same experiment by titrating unlabeled troponin C (in the absence of troponin I), we observed a transfer efficiency shift in the presence of 2 mM CaCl_2_ starting from concentrations of 100 nM troponin C; however, we did not see binding in the absence of calcium (**Fig. S3**). By analyzing the respective fractions of bound and unbound populations, we estimated a K_D_ of troponin T for troponin C of 1200 + 60 nM. This result is reasonable, accounting for the fact that troponin I and T share an interface, whereas troponins T and C share fewer contacts within the troponin complex (63).

Interestingly, when titrating increasing concentration of equimolar troponins I and C to labeled **TnT_153,213_**, we observed a small yet clearly resolvable shift of transfer efficiencies to even lower values, indicating a further expansion of the troponin-T linker upon formation of the complete troponin complex compared to troponin T with troponin I in isolation. This implies that addition of troponin C alters the conformation of troponin I, leading to increased binding to troponin T, even though troponin C is not directly involved in forming the interaction interface betweens troponins T and I,. Since troponins I and C alter the troponin-T linker conformation, we further analyzed the smFRET data as we increased the concentration of 1:1 TnI:TnC to evaluate if there is an impact on the apparent affinity for troponin I. The apparent affinity for the complex formation in presence of CaCl_2_ is of 0.13 ± 0.02 nM, which is greater than the affinity of troponin I alone binding to troponin T. This result suggests a positive cooperative effect for troponin T binding to troponin I, mediated by troponin C binding to troponin I.

#### The linker remains flexible and dynamic in the troponin complex

We examined the troponin-T linker in the context of the full troponin complex. Labeled troponin T was mixed with troponins I and C to form the full complex. We computed the corresponding root-mean-square interdye distance of 53 ± 1 Å in presence of CaCl_2_. This is slightly expanded compared to the distance measured for troponin T in isolation.

Next, we wanted to test whether the troponin-T linker remains flexible and dynamic in the context of the full troponin complex. To this end, we compared the lifetime of the donor (in the presence of acceptor) τ*_*DA*_* against the mean transfer efficiency determined by counting the number of photons. If the intrachain dynamics are much faster than interphoton times, the lifetimes are related to the mean transfer efficiency according to **Eq. 2**:

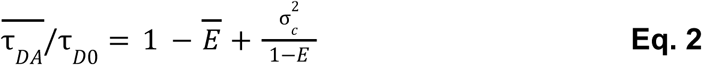

Where 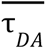 is the mean lifetime associated with the donor-acceptor subpopulation, τ_*D*0_ is the donor in absence of acceptor, and σ*_*c*_* is related to the width of the distribution sampled by the labeled positions (64) (further details in **Supplementary Information, Eq. S7**). Consequently, if the dyes flanked a rigid distance, σ_*c*_ would be equal to zero and the equation would reduce to the classical linear dependence of lifetime and energy transfer that is usually used to interpret FRET measurements. However, if the dyes flank a dynamic region that samples a large conformational space, such as an IDR, σ_*c*_ becomes nonzero, and the data would deviate from the classical linear dependence between lifetime and energy transfer (64).

We measured both the mean lifetime and transfer efficiency for troponin T in isolation and in the context of the full troponin complex (**Fig. 3**). We plotted the theoretical lifetime of the donor versus transfer efficiency for both completely rigid and completely dynamic conformations of the linker (i.e., assuming the region samples the Gaussian chain distribution described by **Eq. S8**). We found both in isolation and in the context of the full troponin complex, that the troponin-T linker shows excellent agreement with the theoretical curve for a dynamic Gaussian chain. This demonstrates that the troponin-T linker is dynamic both in isolation and within the full troponin complex.

**Figure 3:**
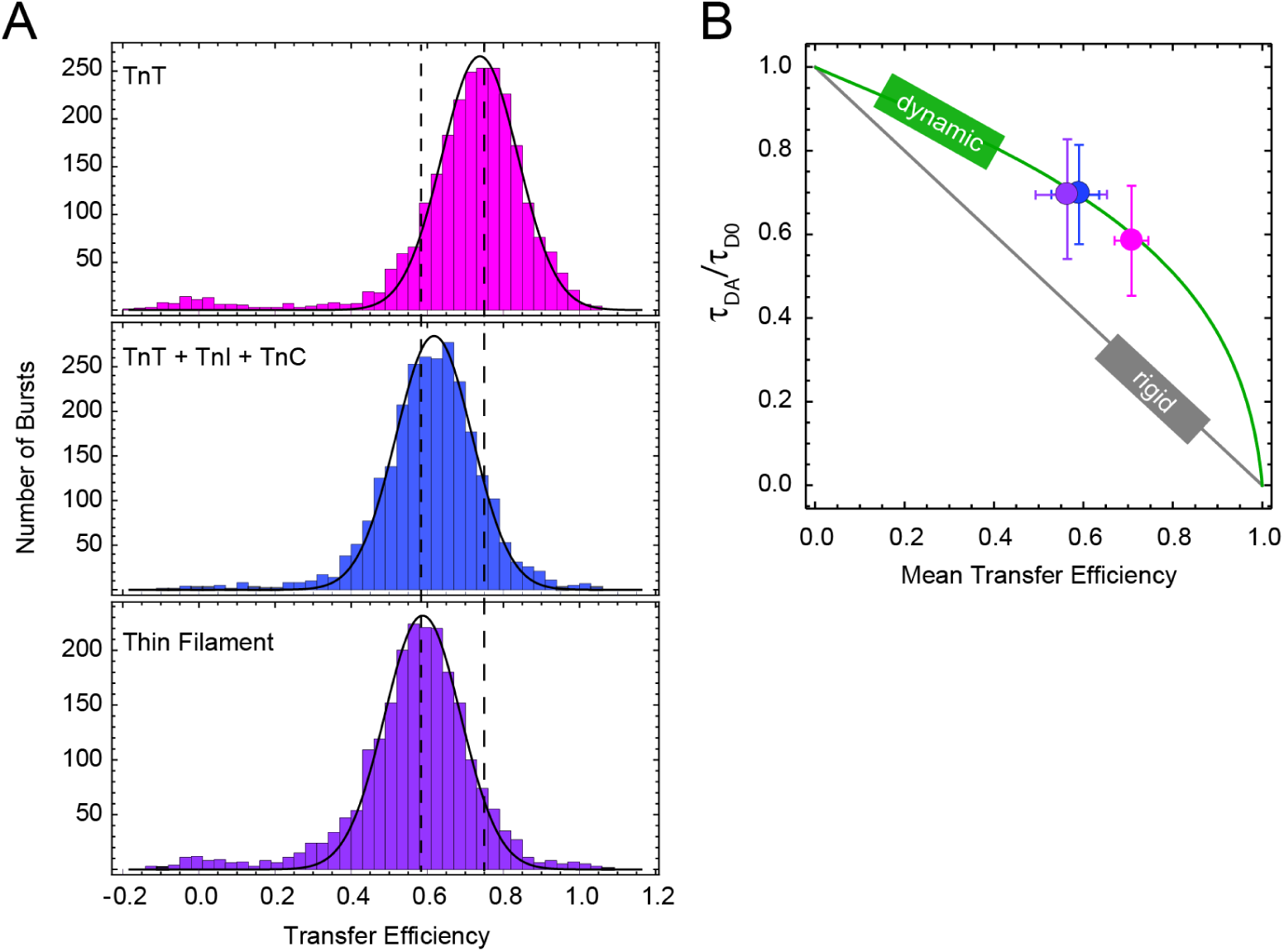
Troponin-T (TnT) linker conformations and dynamics within the troponin complex and thin filament. **A.** Distribution of transfer efficiencies for TnT in isolation (magenta), when part of the troponin complex (blue), and when the troponin complex is bound to the fully regulated thin filament (purple). Vertical lines are guides for the eyes based on the mean transfer efficiencies of TnT in isolation and bound to the thin filament. Troponin I (TnI) and troponin C (TnC) are shown. Corresponding titrations confirming the reported histograms were collected under saturation conditions are shown in **Fig. S3 and S6**. **B**. Normalized lifetime *vs.* transfer efficiency plot for TnT in isolation (magenta), within the troponin complex (blue), and when the troponin complex is bound to the fully regulated thin filament (purple). Values are mean ± standard deviation from independent replicates (at least two measurements). Gray line represents the theoretical static limit of a rigid distance, while the green line describes the corresponding results for a dynamic chain sampling distances according to a Gaussian chain model (see Eq. 2). Data falls on the dynamic curve for all conditions. All experiments performed in 50 mM HEPES, 25 mM KCl, 4 mM MgCl_2_, 2 mM CaCl_2_.

### The troponin-T linker remains dynamic as part of the thin filament

Having established that the troponin-T linker is dynamic and disordered in the context of the full troponin complex, we investigated whether the linker remains dynamic in the context of the fully regulated thin filament. The fully regulated thin filament, composed of actin, troponin, and tropomyosin, is sufficient to recapitulate calcium-dependent regulation of myosin *in vitro* (**Fig. 4**). Using conditions that are sufficient for thin filament assembly and calcium-based regulation *in vitro*, we reconstituted thin filaments containing labeled **TnT_153,213_**.

**Figure 4:**
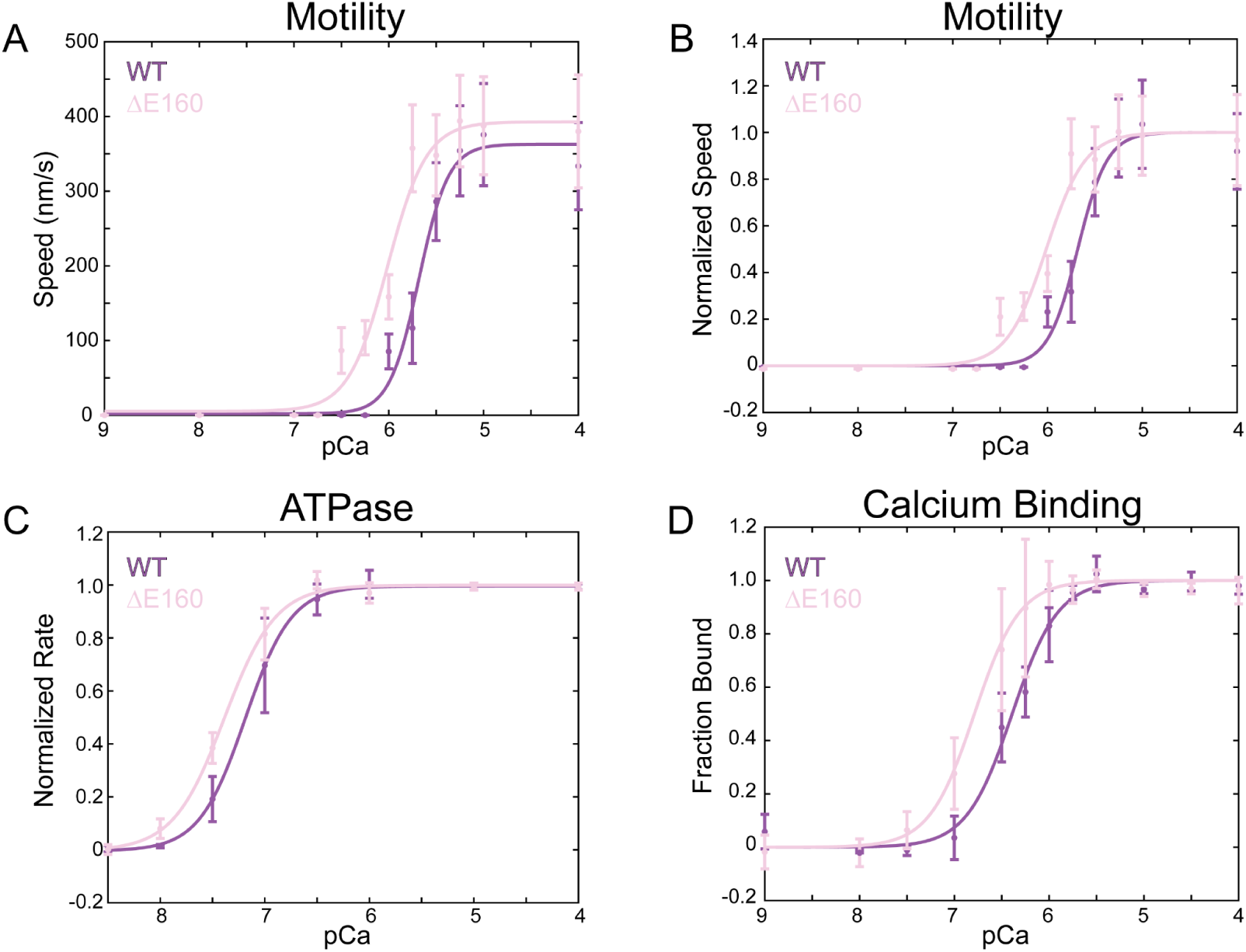
The ΔE160 mutation in troponin T affects thin filament regulation and molecular contractility. **A-B.** *In vitro* motility assays showing the speed of thin filament translocation over a bed of myosin as a function of calcium. Shown are the **A.** un-normalized and **B**. normalized speeds. Data are from N=3 separate days of experiments. The mean and standard deviations are shown. The maximal speed (WT: 360 ± 10 nm/s, ΔE160: 390 ± 10 nm/s, P < 0.01) and the pCa_50_ are increased for the mutant (WT: 5.7 ± 0.1, ΔE160: 6.0 ± 0.1, P < 0.05). The speed is increased for ΔE160 for pCa<6.25 (P < 0.05), except at pCa 5. **C.** The normalized ATPase rate as a function of calcium. Data are from N=5 WT and N=4 ΔE160 curves. The mean and standard deviations are shown. The pCa_50_ is increased for the mutant (WT: 7.2 ± 0.1, ΔE160: 7.4 ± 0.1, P < 0.05). **D.** The fraction of calcium bound to troponin C as a function of calcium measured using IAANS fluorescence. Data are from N=3 WT and N=5 ΔE160 curves. The mean and standard deviations are shown. The pCa_50_ is increased for the mutant (WT: 6.4 ± 0.1, ΔE160: 6.7 ± 0.2, P < 0.05).

We first demonstrated that thin filaments containing the labeled **TnT_153,213_**show slower diffusion through the confocal volume compared to isolated troponin in solution, consistent with the fact that the troponin is bound to the thin filament (**Fig. S4**). Next, we measured the corresponding transfer efficiency distributions for the linker region in the context of the fully regulated thin filament (**Fig. 3**). We observed a single FRET population with a mean transfer efficiency of 0.59 ± 0.02 that is further shifted toward lower values of transfer efficiency compared to isolated troponin T or the troponin complex in solution. These data suggest an additional expansion of the linker when the troponin complex binds to the thin filament. We then examined whether the linker region remains dynamic in the context of the fully regulated thin filament by measuring the relationship between lifetime and transfer efficiency as described above (**Fig. 3**). We found that the linker remains highly flexible in the thin filament and adopts a root-mean-square interdye distance of 55 ± 1 Å. Taken together, our data clearly demonstrate that the troponin-T linker region is disordered and dynamic in isolation, in the troponin complex, and in the context of the fully regulated thin filament.

There are two troponin complexes for every seven actin subunits, and recent cryo-EM studies have demonstrated that these two troponins have different conformations on opposite sides of the actin (7–11). While the troponin-T linker is not resolved for either troponin, the distance from the troponin core complex to the tropomyosin-troponin T overlap region is different for the two troponins. As such, one might have expected to see two different linker distances corresponding to two different troponin conformations; however, we only observe a single peak, consistent with a single population of rapidly interconverting conformations. It is important to note that, even in the context of the thin filament, it is unlikely that we are measuring more than one labeled troponin T molecule at a time, as evidenced by both the stoichiometry ratio measured by PIE and the total number of photons per burst.

We hypothesized that the second population of troponin linker distances might not be observed due to potential differences in affinity of troponin for the two binding sites. To test this, we used crowding agents to increase the local concentration of troponin and potentially reveal a different troponin binding site on the thin filament with a different binding affinity. Indeed, when increasing the volume fraction of the crowder PEG 600, we observe the appearance of a second population with labeling stoichiometry ratio 0.5 (**Fig. 5 A-B**). This new population is characterized by a mean transfer efficiency of 0.29 ± 0.02, and it also falls on the dynamic line, consistent with the notion that this second conformation is also disordered and dynamic. Conversion of the mean transfer efficiency to distance results in a value of 8.6 ± 0.4 nm.

**Figure 5.**
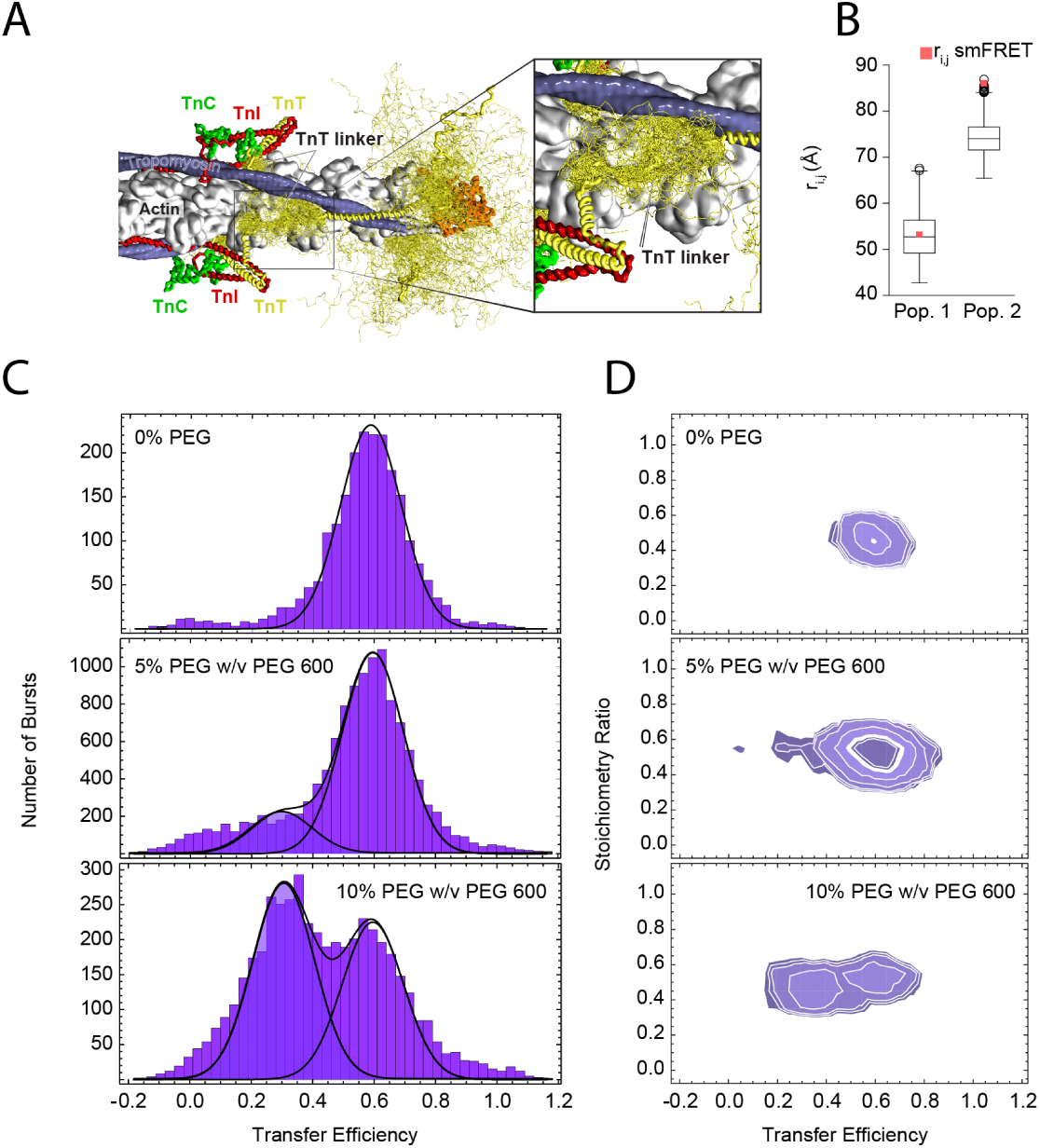
Troponin-T (TnT) linker conformations in the thin filament with crowders. **A.** Simulations highlighting flexibility and extension of the TnT linker (yellow) when in complex with the thin filament comprised of TnC (green), TnI (red), actin (grey), and tropomyosin (blue). **B.** Root mean square inter-dye distances computed from simulations with the single molecule experimental data plotted as an orange square. **C**. Distribution of transfer efficiencies for TnT when the troponin complex is bound to the fully regulated thin filament in buffer (top), with the addition of 5% w/v PEG 600 (middle) and 10% w/v PEG 600 (bottom). **D.** Representative 2D plot of Transfer Efficiency vs labeling stoichiometry ratio for each condition shown in **C**. The slight shift in the mean stoichiometry across the two populations in 10% PEG 600 is due to changes in brightness between the two states (possibly due to quenching) and is not assigned to changes in labeling stoichiometry, which should be 0.33 for 1 donor and 2 acceptor or 0.25 for 1 donor and 3 acceptors. We exclude the case of 2 donors and 2 acceptors because the total number of photons per burst remains similar between the two observed populations.

To assess whether these two populations are compatible with our current understanding of the thin filament structure, we constructed a full thin filament model (15 actin chains, 8 tropomyosin chains, 2 TnT chains, 2 TnI chains, and 2 TnC chains) based on the Ca^2+^-free cryo-EM structure (PDB:6KN7). Using this model, we performed coarse-grained simulations of the fully regulated thin filament with the TnT linkers and N-termini reconstructed (**Fig. 5C**). From these simulations, the average inter-residue distances for dye positions from the two troponins are 5.3 nm (in excellent agreement with the distance obtained in the absence of crowder, 5.4 nm) and 7.4 nm (**Fig. 5D**). While this second population predicted computationally is more compact than the population observed experimentally under crowding conditions, this may reflect constraints imposed by the structural model. Taken together, our results demonstrate the ability to resolve two different, yet still dynamic conformations of the linker on the thin filament, and suggest that the two troponin complexes on opposite sides of the thin filament have different binding affinities.

### The ΔE160 mutation in the linker region affects thin filament activation

Given that the troponin-T linker remains dynamic and disordered both in the troponin complex and the fully regulated thin filament (**Fig. S7**), we asked how pathogenic mutations within the linker region impact troponin function. We focused on the ΔE160 mutation located in the linker region that has been extensively studied in multiple model systems and clearly demonstrated to be pathogenic, causing hypertrophic cardiomyopathy (13, 15, 27, 65–72).

To demonstrate the functional effects of the ΔE160 mutation using the same protein components and buffer conditions used in the single-molecule experiments, we expressed the human WT and ΔE160 troponin T proteins and reconstituted them into troponin complexes (**Fig. 4**). We then generated functional regulated thin filaments composed of human troponin and tropomyosin and porcine cardiac actin, which is identical to human cardiac actin. Moreover, we conducted our experiments using porcine cardiac myosin, which has biophysical properties that are indistinguishable from human cardiac myosin (31, 73, 74).

To examine the effects of ΔE160 on contractility, we used the in vitro motility assay in which fluorescently-labeled regulated thin filaments are translocated over a bed of myosin in the presence of ATP and calcium (75). Consistent with previous studies (70–72), we observed that while both WT and ΔE160 thin filaments show calcium-based regulation, ΔE160 shows increased activation at lower calcium concentrations compared to the WT (**Fig. 4**). Data were fitted with the Hill equation, and we observed a shift in the pCa_50_ towards activation at lower levels of calcium for ΔE160 as well as an increase in the maximal speed of movement at saturating calcium concentrations. This means that ΔE160 requires less calcium to reach the same level of thin filament activation, consistent with molecular hypercontractility.

Next, we examined the effects of ΔE160 troponin T on thin filament regulation by measuring the myosin ATPase rate in the presence of regulated thin filaments as a function of calcium. Consistent with previous studies (70–72), both the WT and mutant proteins showed low ATPase activity at low calcium, and high activity at saturating concentrations of calcium (**Fig. 4**) Data were fitted with the Hill equation to extract the calcium concentration at half-maximal activation (pCa_50_). We found that ΔE160 showed a shift in pCa_50_ towards activation at lower calcium concentrations.

Taken together, both the ATPase and in vitro motility experiments demonstrate that the mutation causes increased activation at lower calcium concentrations, consistent with molecular hypercontractility and previous studies of the ΔE160 mutation.

### ΔE160 mutation does not alter conformations but interactions

Having established that the ΔE160 mutation affects the functional properties of the reconstituted thin filament, we examined whether the mutation affects the linker structure, dynamics, and/or troponin T’s interactions with binding partners. Importantly, the ATPase and motility assays demonstrated quantifiable mutational effects with a limited number of 5 protein components. Therefore, we can test the role of each of these components.

First, we examined the properties of ΔE160 troponin T in isolation. We measured the distribution of transfer efficiencies, finding a negligible variation in the mean value compared to wildtype (**Fig. 6**). Lifetime measurements and nsFCS confirmed that the linker of ΔE160 is flexible and dynamic with a relaxation time of 130 ± 20 ns. The addition of KCl reveals a minor but quantifiable difference at higher ionic strength (0.5 M), with the ΔE160 variant expanding more than wild-type (**Fig. 6**). From the mean transfer efficiency, we computed a root-mean-square interdye distance of 40 ± 1 Å. Overall, the comparison of the distribution of transfer efficiencies between wild-type and mutant troponin T indicates a minor change in the conformations and dynamics of the linker.

**Figure 6:**
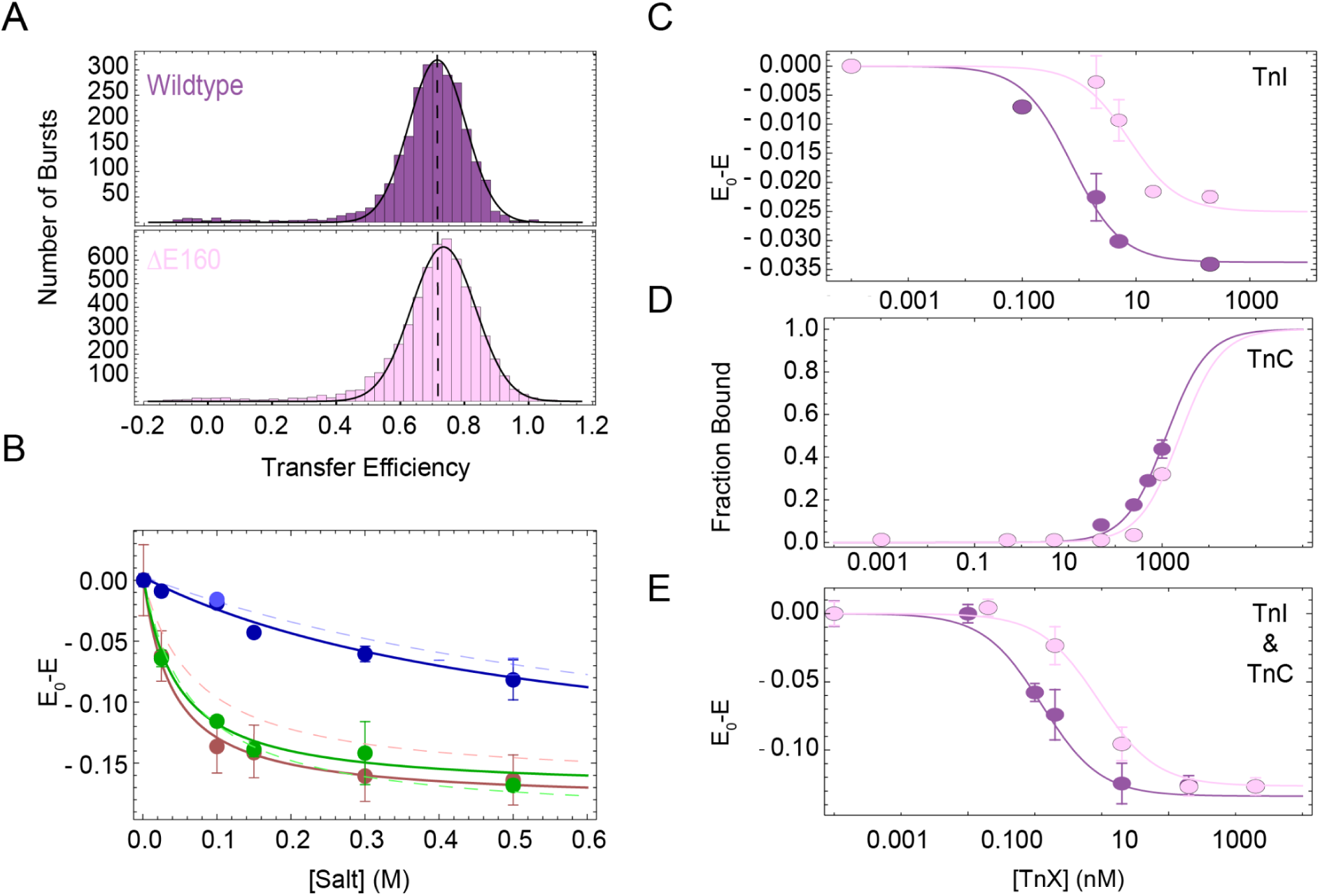
Effects of ΔE160 mutation on conformations, interactions, and salt response. **A.** Comparison of distribution of transfer efficiencies in 50 mM HEPES, 25 mM KCl, 4 mM MgCl_2_, and 2 mM CaCl_2_ for **TnT_153,213_** wild-type (purple) and ΔE160 (light pink). **B.** Mean transfer efficiency of **TnT_153,213_** as a function of increasing concentration of salt (KCl in blue, CaCl_2_ in green, MgCl_2_ in pink), compared to WT (dashed lines). **C-E.** Comparison of binding curves of troponins T (TnT) and I (TnI) (**C**), TnT and troponin C (TnC) (**D**), and TnT and 1:1 TnI:TnC concentration (**E**) for wild-type (purple) and ΔE160 (light Pink) **TnT_153,213_**. All errors are standard deviations determined from independent replicates of the same sample (at least two measurements). Representative titrations in **Fig. S3,S5-6**.

We then investigated whether the mutation alters the interactions of the linker with the other troponin components. Interestingly, we found that binding curves for troponin I are shifted toward higher values, with K_D_ values of 3.2 ± 1.4 nM in absence of calcium and of 7.4 ± 3.6 nM in presence of CaCl_2_, differing by more than a factor of 10 compared to WT (**Fig. 5 and S5**). This suggests that the ΔE160 mutation in the linker affects intermolecular interactions with other troponin subunits. Moreover, similar to WT, when titrating troponin C, we only observe a shift towards more expanded transfer efficiencies in the presence of CaCl_2_, with a K_D_ value of 2400 ± 300 nM. When we mix equimolar concentrations of TnI and TnC, we found that the binding curve is shifted toward a higher value, with a K_D_ value of 2.6 ± 0.5 nM in presence of calcium, differing by more than a factor of 10 compared to WT (**Fig. 6 and S5**).

Since the ΔE160 mutation appears to affect binding interactions with troponins I and C, we further investigated whether the ΔE160 mutation in the linker region affects the binding of calcium to troponin C, located in the core of the troponin complex. We used a modified troponin C construct that was labeled with IAANS dye to report calcium binding (48). We then reconstituted this troponin C into full regulated thin filaments containing actin, tropomyosin, and the troponin complex. This is a well-established approach for spectroscopically measuring calcium binding to troponin C. We performed a titration curve with increasing concentrations of calcium. We found that both the WT and ΔE160 troponins bind calcium, leading to alterations in the fluorescence. Consistent with previous studies (72), we observed that the ΔE160 shows increased calcium binding to troponin C compared to the WT (**Fig. 4**). This result demonstrates that in the context of the fully regulated thin filament, ΔE160 can affect calcium binding to troponin C. This is consistent with our binding measurements, showing that despite being spatially separated from the troponins I and C binding sites, the ΔE160 mutation can allosterically affect the conformation and functional properties of binding partners.

## DISCUSSION

Troponin plays a central role in the regulation of calcium-dependent interactions between myosin and the thin filament, and it is well established that mutations in cardiac troponin can disrupt muscle contraction and lead to cardiomyopathies (76). As such, the structure-function relationship of the thin filament has been an active area of research for decades. Excellent structural studies using crystallography and electron microscopy have resolved key regions of the thin filament, both in isolated proteins and in the context of the fully regulated thin filament; however, several key regions of troponin have remained unresolved, including the troponin-T linker region (7–11, 63).

Here, we applied computational and biophysical approaches to probe the troponin-T linker region. We demonstrate that this region is a dynamic IDR in the context of isolated troponin T, as part of the troponin complex, and in the fully regulated thin filament. Moreover, we demonstrate that a mutation causing a single amino acid deletion in this region associated with hypertrophic cardiomyopathy, ΔE160, primarily affects the interactions with other subunits in the troponin complex rather than the conformational ensemble or dynamics of the linker region. These results have important implications for our understanding of both structure-function relationships of muscle proteins and the pathogenic mechanisms that can drive cardiomyopathies.

### The troponin-T linker region is dynamic and disordered

Here, we used single-molecule techniques to experimentally test the computational prediction that the troponin-T linker would be disordered. We show that in the context of the isolated troponin-T complex, this region displays several hallmarks of an intrinsically disordered region, including: i) this region samples many configurations on a 100 nanosecond time scale, ii) the linker region expands monotonically upon the addition of GdmCl, suggesting the absence of persistent secondary structure over a substantial number of residues, and iii) the root-mean-square distance between the fluorescence probes is 4.60 ± 0.05 nm, consistent with the results from molecular dynamics simulations in which the linker is fully disordered. We see similar results with the full troponin complex. Taken together, our data clearly demonstrate that the troponin-T linker region is disordered in solution.

While disorder predictors all indicate this region should be disordered, recent work has called into question whether highly-charged blocky sequences, such as the linker sequence, truly exist as disordered regions and may instead exist as alpha helices (18, 77–79). While we cannot extrapolate beyond our data, at least for the troponin-T linker, we see no evidence of helicity or changes in IDR dynamics under the range of conditions measured. This is consistent with the cryo-EM structure obtained under both low and high calcium conditions. This notwithstanding, we cannot rule out the possibility that transient structure is acquired at some point during thin filament cycles or regulation, but the most parsimonious conclusion from our data thus far is that the troponin-T linker is constitutively disordered.

We confirmed that the sequence of the troponin-T linker region has a polyampholytic response, where addition of salt leads to screening of attractive positive and negative interactions and consequent expansion of the conformational ensemble. Intriguingly, the degree of expansion in the presence of divalent ions is stronger than with monovalent ions, beyond the simple expectations due to the differences in ionic strength. As suggested by polymer theories (80), the measured conformations must arise because of the specific role of counterion condensation. Condensation of monovalent ions can result in neutralization of oppositely charged residues, whereas condensation of divalent counterions can lead to charge reversal (because of the excess of charge of the counterion) or bridging (connecting two charged residues within the protein) (80). While we did not observe a significant change in the linker response between either calcium or magnesium divalent salts, our observation clearly points to an important difference in the linker response that depends on the nature of the ions in solution. Therefore, conformations explored by the linker are not only influenced by the sequence of the protein, but also by the condensation of specific ions. This may partially explain the temperature insensitivity of the linker that we observed, since counterion release with increasing temperature may be compensated by charge reversal. However, we cannot exclude that thermal denaturation of the surrounding domain also compensates for the expected compaction, though this would be an unexpected result since a reduction in the excluded volume of the folded domain usually results in further compaction of the chain, not expansion.

We also applied our tools to examine the conformation and dynamics of the troponin-T linker in the context of the fully regulated thin filament. We find that the FRET efficiency between the probes is well described by a single, shot-noise limited Gaussian with a mean distance between the FRET probes of 55 ± 1 Å in the presence of calcium. This distance is similar to the distance predicted from our simulations and the mean distance previously measured using ensemble FRET lifetimes (81).

Recent structural studies of the thin filament have demonstrated that the linker regions on opposite sides of the thin filament adopt different conformations. We were able to observe a second population of distances, but only in the presence of crowding agents. Conversion of the mean transfer efficiency for this second population to distance results in a value of 86 ± 4 Å, which is in general agreement with distances predicted from simulations. Since this population only emerges in the presence of crowding agents, this suggests that the two conformations of the troponin linker have different affinities for the thin filament. Under the low concentrations of troponin used for smFRET experiments, we only observe the less extended conformation, consistent with a higher affinity of that conformation for the thin filament. It is worthwhile to note that in solution (unbound), the linker has a mean length *R*_*unbound*_ 46 ± 5 Å. This is similar to the conformations in the troponin complex (*R*_*tc*_ = 53 ± 1 Å) and to the less extended population on the thin filament (*R_*tf*_* = 55 ± 1 Å). As such, it is possible that the linker with the less extended conformation has a higher affinity for the thin filament since it does not require as much extension of the linker upon binding to the thin filament compared to the second, more extended conformation. Assuming the linker behaves as an entropic spring, we calculated the free energy cost for the conformational change in the linker upon binding to the thin filament in the two configurations 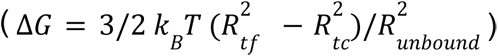, resulting in a Δ*G* of 0.2 ± 0.6 *k*_*B*_*T* and 3.3 ± 1.2 *k*_*B*_*T* for the closed and expanded configurations, respectively. If the cost is transferred completely to the binding affinity, this would correspond to a change of a factor of 30 between the affinities of the two sites.

Aside from measuring the mean distances explored by the linker, our approach also allows us to measure the dynamics of the linker region by accounting for how FRET affects the fluorescence lifetime of the donor while the probes dynamically sample the distance distribution (see Supplementary Information, in particular, Eq. **S5-S8**). Our lifetime measurements demonstrate that the troponin-T linker region is highly dynamic within the context of fully reconstituted human troponin complex and the fully regulated thin filament (**Fig. 3**). Taken together, our results clearly demonstrate that the troponin-T linker region is disordered and dynamic, even in the context of the fully regulated thin filament. As such, our results potentially explain why this region has been elusive to structural biology techniques. Moreover, they demonstrate the importance of accounting for the linker disorder when modeling the structure of the thin filament.

### The ΔE160 mutation affects contractility through altered intermolecular interactions

The ΔE160 mutation in the linker region of troponin T is a well-established mutation that causes hypertrophic cardiomyopathy in patients. This mutation has been investigated in multiple model systems ranging from purified proteins to animal models, and it has been shown to cause increased contractility across scales (13, 15, 27, 65–72). Consistent with these previous studies, we demonstrate that the ΔE160 mutation causes a shift towards submaximal calcium activation in both the in vitro motility assay and the calcium-dependent thin-filament activated ATPase rate. Similar shifts have previously been observed using reconstituted proteins and isolated muscle fibers (70–72). Moreover, similar shifts towards submaximal calcium activation have been previously observed with other troponin T mutations associated with hypertrophic cardiomyopathy, and this shift has been proposed to cause increased contractility in response to a calcium transient (12, 30, 82–84). As such, our results clearly demonstrate that we can recapitulate functional changes in molecular-based contractility using our simplified system.

Multiple structural models have been put forward to explain the effects of mutations in the linker assuming that the linker behaves like other well-defined, folded proteins (67, 81, 85); however, some of these assumptions underlying these models would not apply to intrinsically disordered proteins that rapidly explore diverse conformational ensembles. We investigated whether the ΔE160 mutation affects the structure and/or dynamics of the linker region. Using both isolated and complexed ΔE160 troponin-T, we observe that the linker region remains dynamic and disordered. When compared to the WT protein, we observe that the mean transfer efficiency is essentially unchanged in the ΔE160 protein. Moreover, the ns-FCS measurements show that the dynamics of the linker are unchanged by the ΔE160 mutation, reporting a relaxation time of 130 ± 20 ns for the ΔE160 mutant compared to 120 ± 10 ns for the WT. When reconstituted into the fully regulated thin filament, we observed that the linker region is similarly dynamic with a similar mean distance between the probes as the WT. Taken together, our results demonstrate that the ΔE160 mutation does not significantly change the conformational dynamics of the linker region.

While the ΔE160 mutation did not affect the structural dynamics of the linker region, it had a clear effect on the interactions of troponin T with troponins I and C. The apparent binding affinity of troponin T for troponins I and C is ∼ 10-fold weaker for the mutant compared to the WT, with a K_D_ value of 0.9 ± 0.2 nM. This result is seen in solution, outside of the context of the fully regulated thin filament, demonstrating that the ΔE160 mutation in the disordered linker leads to allosteric changes that affect the intermolecular interactions between troponin subunits. Consistent with this notion, we measured the equilibrium binding of calcium to the troponin C in the context of the fully regulated thin filament. Our data clearly demonstrate that the ΔE160 mutation in the disordered troponin-T linker increases the binding of calcium to the troponin core complex, despite the spatial separation between these structures. Moreover, the shift towards submaximal calcium binding is similar to the shifts towards increased activity at submaximal calcium activation seen in the functional ATPase and in vitro motility assays. As such, we propose the ΔE160 mutation exerts its effects primarily through allosteric effects on the interactions between troponin subunits rather than through direct effects on the structural dynamics of the linker. While additional studies are needed to resolve the details of this coupling, we propose that other mutations in the disordered linker could also exert their effects through similar mechanisms.

#### Conclusions

Here, we demonstrate that the troponin-T linker region is dynamic and disordered both in isolation and in the context of the fully regulated thin filament. As such, modeling this region requires consideration of how intrinsically disordered regions behave differently from structured regions. We propose that rather than thinking of how changes in sequence affect the structure of the linker region, it is more informative to consider how changes in sequence alter the conformational ensembles and dynamics of the IDR, leading to altered molecular function.

## ACKNOWLEDGEMENTS

The authors would also like to acknowledge the financial support provided by the National Institutes of Health (R01 HL141086 to M.J.G., R01 AG062837 to A.S. and AI163142 to A.S and A.S.H.), the Children’s Discovery Institute of Washington University and St. Louis Children’s Hospital (PM-LI-2019-829 to M.J.G.), and the Longer Life Foundation (an RGA/Washington University Collaboration, to A.S.H.). J.C. is supported by the W.M. Keck Foundation with a W.M. Keck Postdoctoral Fellowship.

## AUTHOR CONTRIBUTIONS

Conception, funding, and oversight by M.J.G., A.S.H., and A.S. L.G. and A.E.G. generated protein constructs and performed functional experiments. J.C. labeled the protein constructs, performed, and analyzed the single-molecule experiments. A.S. and M.D.S.B. oversaw single-molecule experiments and analysis and developed analytical tools. A.S.H and R.J.E. designed, performed, and analyzed simulations. All authors contributed to the analysis of the data. The first draft was written by M.J.G., A.S., and A.S.H.. All authors contributed to the writing and/or editing of the manuscript.

