## Supplemental Materials for "Structural dynamics of the intrinsically disordered linker region of cardiac troponin T"

$$E = n_A/(n_A + n_D) \qquad \textbf{Eq. S1}$$

where $n_A$ and $n_D$ are the donor and acceptor counts per burst corrected for background, crosstalk, excitation, and variation in quantum yield. For the typical case of a fixed rigid distance $r$, the lifetime of the donor in the presence of an acceptor is given by

$$\tau_{DA} = \tau_{D0}(r)(1 - E(r)) \qquad \textbf{Eq. S2}$$

where

$$E(r) = \frac{R_0^6}{R_0^6 + r^6} \qquad \textbf{Eq. S3}$$

However, for the case of disordered regions, the protein does not adopt a single conformation, but many conformations, and this requires properly accounting for the underlying distribution in the estimation (or conversion) of the mean transfer efficiency, the fit of the lifetime decay, and the conversion of the mean lifetime to distance.

In this scenario the mean transfer efficiency is given by

$$\overline{E} = \int E(r)P(r)dr \qquad \textbf{Eq. S4}$$

The time-dependent fluorescence intensity decay of the donor-acceptor population is given by:

$$I(t) = I_0 \int P(r) e^{-t/\tau_{DA}(r)} dr \qquad \textbf{Eq. S5}$$

and the mean lifetime $\overline{\tau_{DA}}$ is given by

$$\overline{\tau_{DA}} = \int t\, I(t)\, dt / \int I(t)\, dt \qquad \textbf{Eq. S6}$$

$\overline{\tau_{DA}}$ can be further related to the mean transfer efficiency $\bar{E}$ using **Eq. 2**, where:

$$\sigma_c^2 = \int E^2(r) P(r) dr - \left( \int E(r) P(r) dr \right)^2 \qquad \textbf{Eq. S7}$$

The parameter $\sigma_c^2$ reflects the broad range of configurations sampled by the chain and predictions can be obtained assuming a Gaussian distribution:

$$P(r) = 4\pi r^2 \left( \frac{3}{2\pi \overline{r^2}} \right)^{3/2} e^{-\frac{3}{2}\frac{r^2}{\overline{r^2}}} \qquad \textbf{Eq. S8}$$

This is an essential element to consider when converting distances based on the lifetime or on the mean transfer efficiency estimated from the single-molecule bursts.

**nsFCS analysis.** Autocorrelation curves of acceptor and donor channels and cross-correlation curves between acceptor and donor channels were computed as described previously (55, 86). Measurements were performed at ~100 pM, and bursts corresponding to the donor-acceptor population were selected to eliminate the contribution of donor-only to the correlation amplitude. The correlation was computed over a time window of 5 μs, and fit to:

$$g_{ij}(\tau) = 1 + \frac{1}{N}(1 - c_{AB}Exp[-(\tau - \tau_0)/\tau_{AB}]) \times$$
$$\times (1 + c_{CD}Exp[-(\tau - \tau_0)/\tau_{CD}])(1 + c_{T}Exp[-(\tau - \tau_0)/\tau_{T}]) \quad \textbf{Eq S9}$$

where *N* is the mean number of molecules in the confocal volume and $i$ and $j$ identify the type of signal (either Acceptor or Donor). The three multiplicative terms describe the contribution to amplitude and timescale of photon antibunching (AB), chain dynamics (CD), and triplet blinking of the dyes (T).

**Binding model.** Binding of ligands $L$ to labeled **$TnT_{153,213}$** was studied by following either the mean value of the transfer efficiency distribution or the fraction of bursts associated with the bound/unbound population (when they can be distinctly resolved). Two different equations were used to extract the corresponding binding dissociation constant $K_D$.

In the first case, titration curves were analyzed according to:

$$\overline{E} - \overline{E_f} = (\overline{E_b} - \overline{E_f})\frac{[L]}{K_D + [L]} \quad \textbf{Eq. S10}$$

In the second case, the titration was analyzed according to:

$$f_b = \frac{[L]}{K_D + [L]} \quad \textbf{Eq. S11}$$

### SUPPLEMENTARY FIGURES.

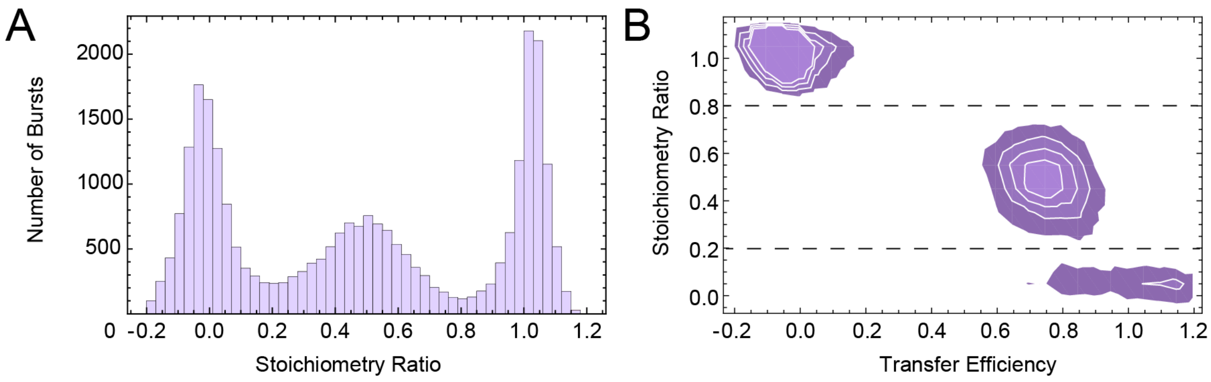


**Supplementary Figure 1. Labeling stoichiometry ratio from pulsed interleaved excitation (PIE): A.** Representative histogram of bursts at given stoichiometry ratios, where a value of one is equal to donor only, a value of zero is equal to acceptor only, and a value of 0.5 indicates molecules with one donor and one acceptor molecule. **B.** Representative 2D plot of Transfer Efficiency vs Stoichiometry ratio. The population at the given transfer efficiency contains only 0.5 stoichiometry ratio.

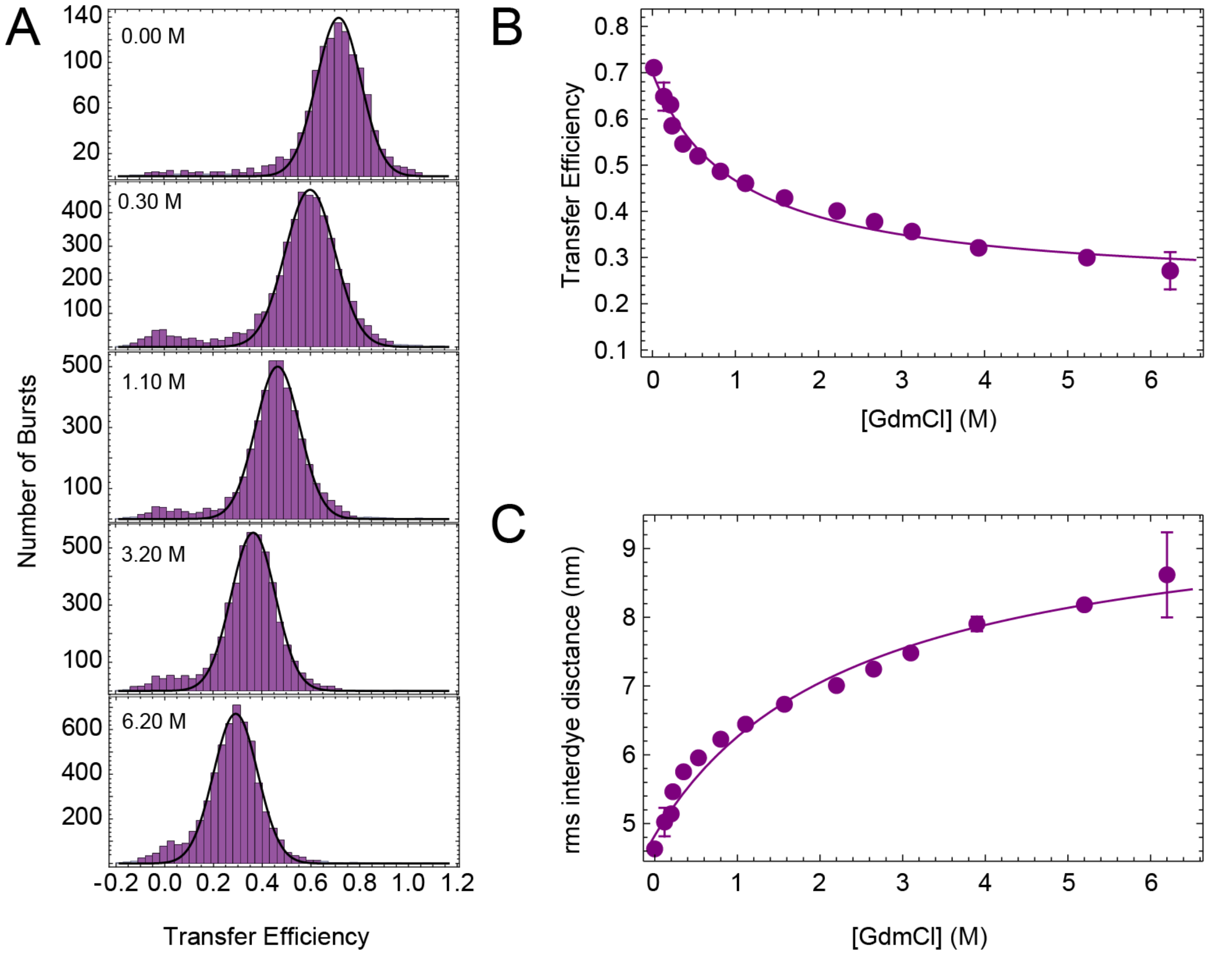


**Supplementary Figure 2. Denaturant titration of the troponin-T (TnT) linker. A.** Representative histograms of the distribution of transfer efficiencies measured for TnT linker as a function of GdmCl concentration. B. Mean transfer efficiency as a function of denaturant concentration. Values are mean ± standard deviation from independent repeats (at least two measurements). Line is a fit to the data based on a Schellman weak binding model. C. Root-mean-squared interdye distance as a function of denaturant concentration. Values are mean ± standard deviation from independent repeats (at least two independent measurements; if error bars are not shown, they are of comparable or smaller size than the dots). Line is a fit to the data based on the Schellman weak binding model.

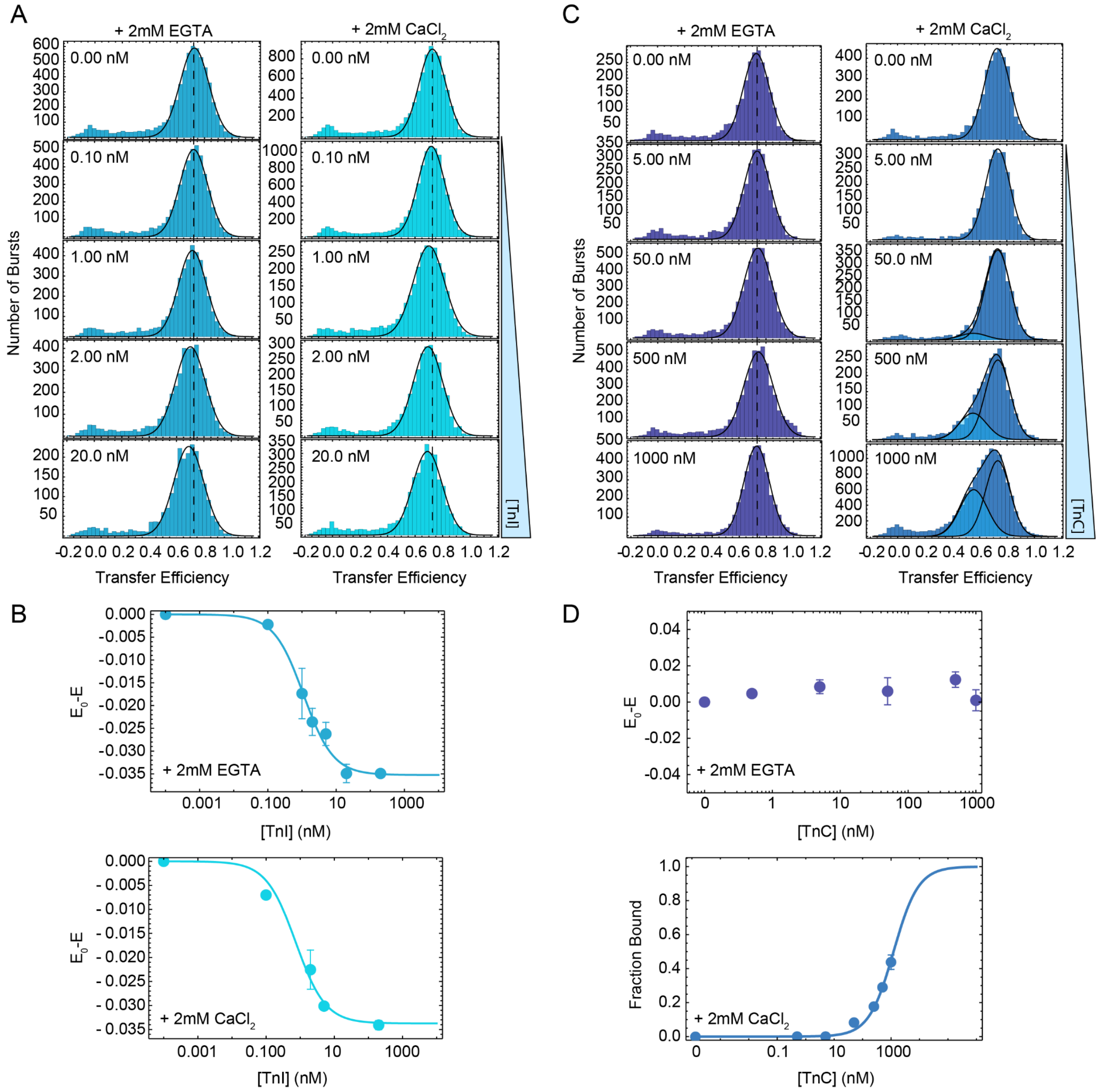


**Supplementary Figure 3. Troponin T binding to troponins I (TnI) and C (TnC) in absence and presence of $CaCl_2$. A.** Representative distribution of transfer efficiency for the **$TnT_{153,213}$** upon interaction with TnI in the presence of EGTA (blue) and divalent ions (cyan). **B.** Mean transfer efficiency shift as a function of concentration and corresponding binding curve assuming a 1:1 stoichiometry binding model. **C.** Representative distribution of transfer efficiency for the **$TnT_{153,213}$** upon interaction with TnC in presence of EGTA (purple) and divalent ions (dark blue). Since a clear shoulder is observed at high concentration of TnC when divalent ions are present, we fit the data with two Gaussian distributions. **D.** Mean transfer efficiency shift as a function of concentration for the EGTA condition. Values are mean and standard deviation of multiple independent repeats. For the divalent ion condition, fraction bound is fitted to a 1:1 stoichiometry binding model for extracting the corresponding binding affinity.

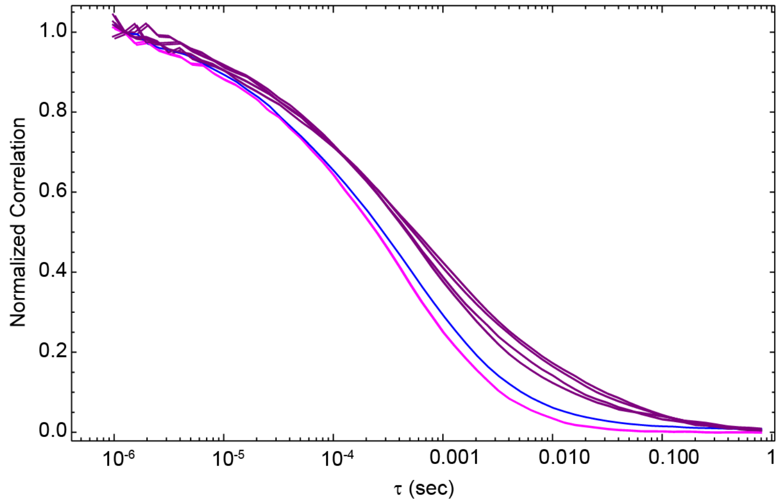


**Supplementary Figure 4. FCS measurements of TnT in isolation, in the troponin complex, and bound to the regulated thin filament.** Representative correlation curves of labeled **$TnT_{153,213}$** in 50 mM HEPES, 25 mM KCl, 4 mM $MgCl_2$, and 2mM $CaCl_2$ for **$TnT_{153,213}$** in isolation (magenta), bound in the troponin complex (blue), and within the thin filament (purple). Note that small fragments of actin are used for the reconstitution of the thin filament in the single-molecule FRET and FCS experiments. Multiple curves are provided to show reproducibility of the results with the thin filament in independent preparations.

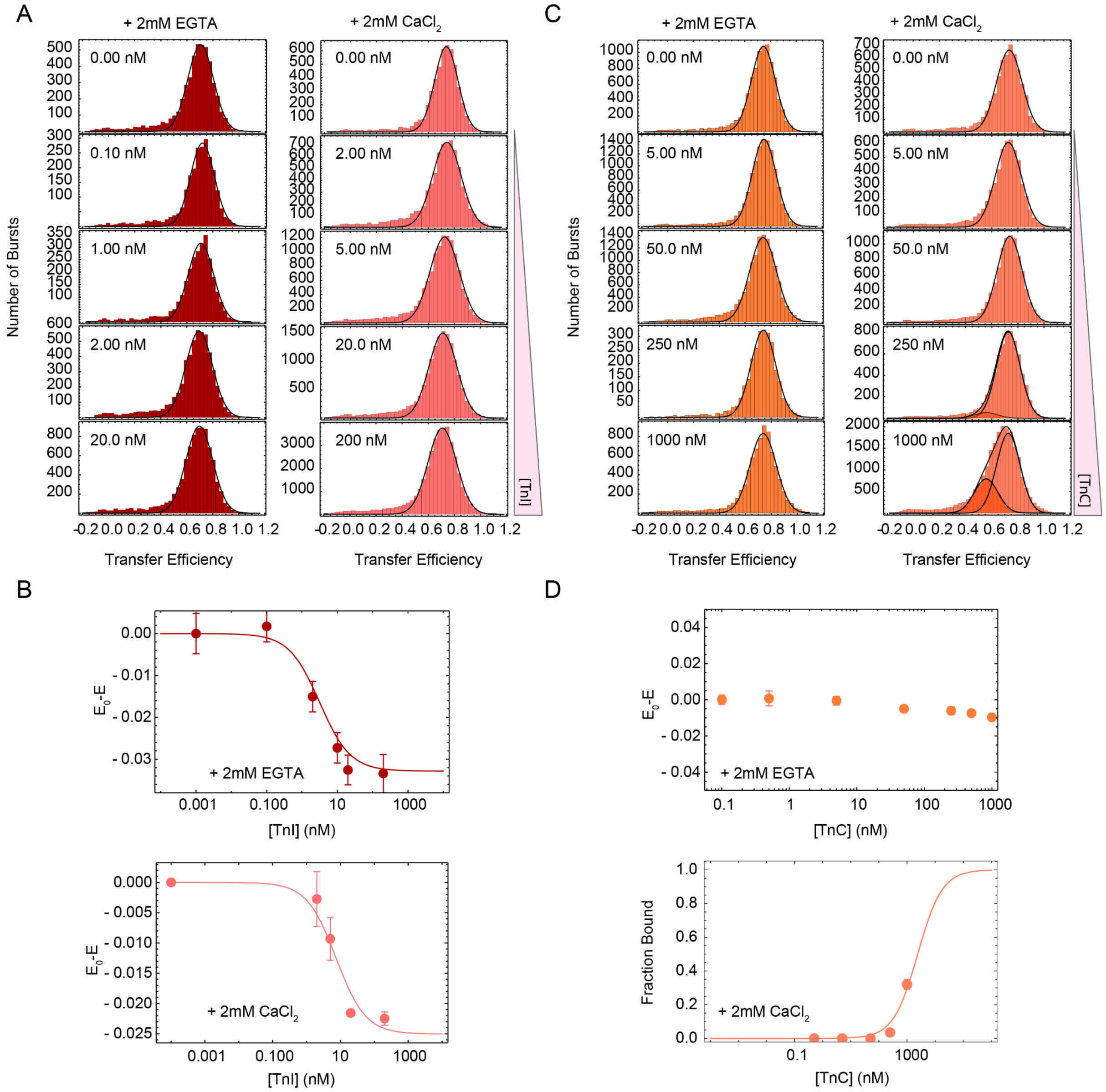


**Supplementary Figure 5. ΔE160 mutant binding to TnI and TnC in absence and presence of $CaCl_2$. A.** Representative distribution of transfer efficiencies for the **$TnT_{153,213}$** upon interaction with TnI in presence of EGTA (red) and $CaCl_2$ (light red). **B.** Mean transfer efficiency shift as a function of concentration and corresponding binding curve assuming a 1:1 stoichiometry binding model. **C.** Representative distribution of transfer efficiencies for the **$TnT_{153,213}$** upon interaction with TnC in the presence of EGTA (orange) and divalent ions (light orange). Since a clear shoulder is observed at high concentration of TnC when divalent ions are present, we fit the data with two Gaussian distributions. **D.** Mean transfer efficiency shift as a function of concentration for the EGTA condition. Values are mean and standard deviation of multiple independent repeats. For the divalent ion condition, fraction bound is fitted to a 1:1 stoichiometry binding model for extracting the corresponding binding affinity.

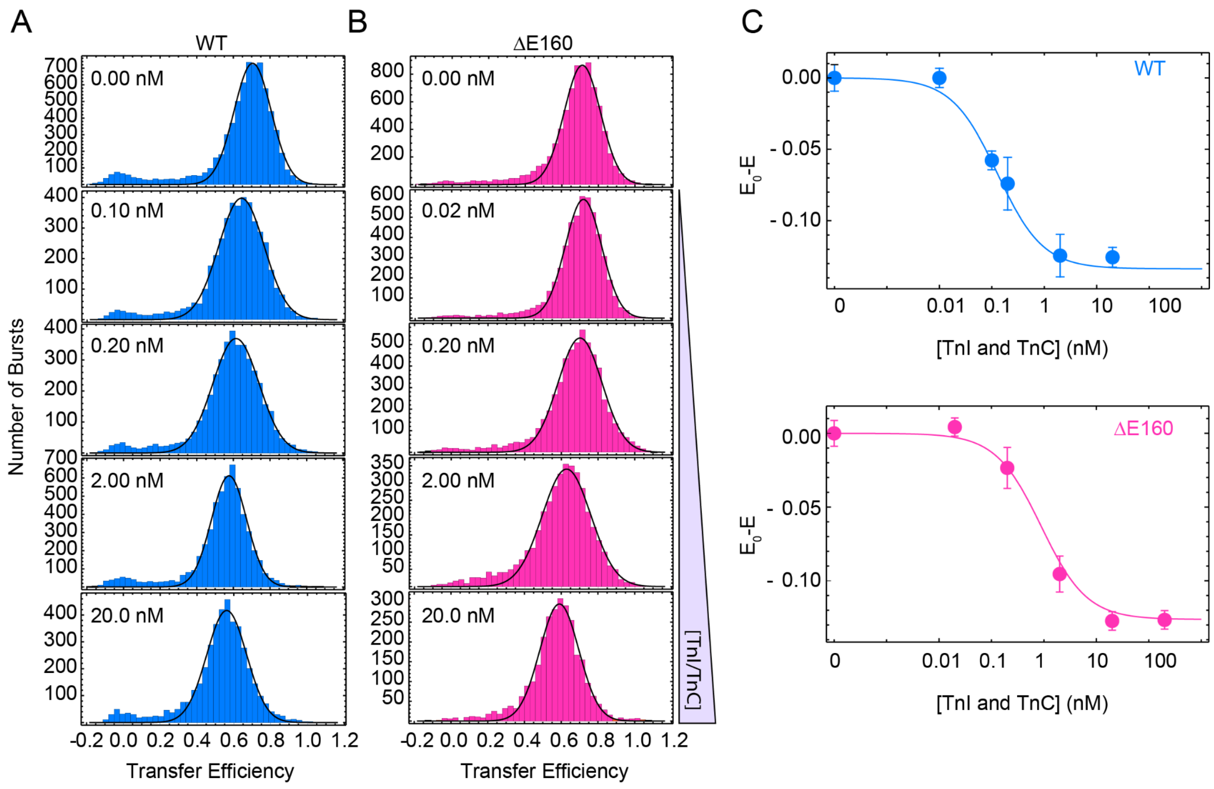


**Supplementary Figure 6. Binding titrations for WT and the ΔE160 mutant to both TnI and TnC in the presence of 2 mM $CaCl_2$. A-B.** Representative titration of transfer efficiency for the **$TnT_{153,213}$** upon interaction with 1:1 TnI:TnC in presence of $CaCl_2$ for wild-type protein (Blue, abbreviated WT) and the ΔE160 mutant (Pink). **C-D.** Mean transfer efficiency shifts as a function of concentration of TnI:TnC for wild-type (blue) and ΔE160 (pink) conditions. Values are mean and standard deviation of multiple independent repeats (at least two). For the divalent ion condition, fraction bound is fitted to a 1:1 stoichiometry binding model for extracting the corresponding binding affinity.

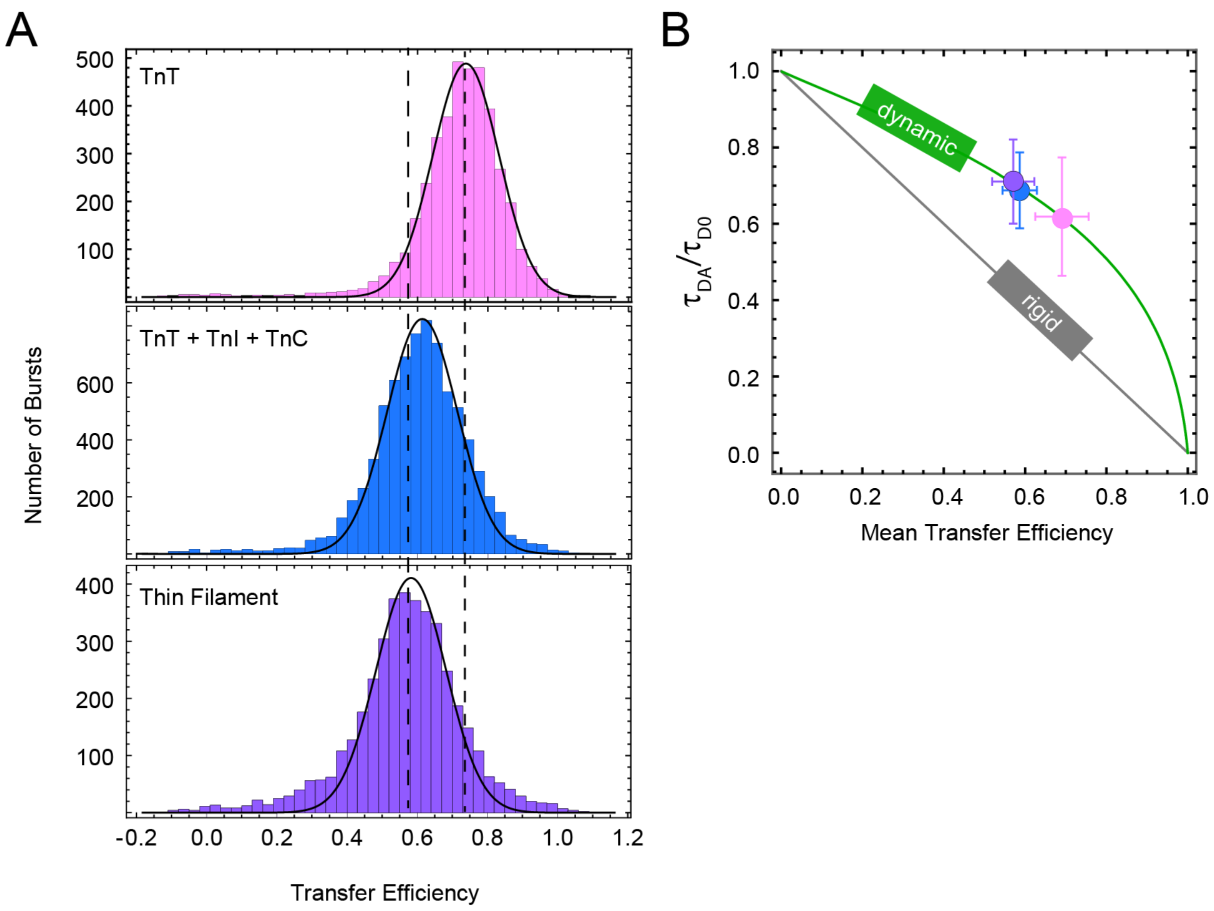


**Figure S7: Troponin-T (TnT) ΔE160 linker conformations and dynamics within the troponin complex and thin filament. A.** Distribution of transfer efficiencies for TnT in isolation (pink), when part of the troponin complex (blue), and when the troponin complex is bound to the fully regulated thin filament (purple). Vertical lines are guides for the eyes based on the mean transfer efficiencies of TnT in isolation and bound to the thin filament. Troponin I (TnI) and troponin C (TnC) are shown. Corresponding titrations confirming the reported histograms are at saturation conditions are shown in **Fig. S5 and S6**. **B.** Normalized lifetime *vs.* transfer efficiency plot for TnT in isolation (pink), within the troponin complex (blue), and when the troponin complex is bound to the fully regulated thin filament (purple). Values are mean ± standard deviation from independent replicates (at least two measurements). Gray line represents the theoretical static limit of a rigid distance, while the green line describes the corresponding results for a dynamic chain sampling distances according to a Gaussian chain model (see Eq. 2). Data fall on the dynamic curve for all conditions. All experiments performed in 50 mM HEPES, 25 mM KCl, 4 mM $MgCl_2$, and 2 mM $CaCl_2$.

### SUPPLEMENTARY TABLES

**Supplementary Table 1.** Sequence of Wildtype (WT) troponin T Protein. Labeling positions are reported as bold and underlined residues.

1 MSDIEEVVEE YEEEEQEEAA VEEQEEAAEE DAEAEAETEE TRAEEDEEEE

51 EAKEAEDGPM EESKPKPRSF MPNLVPPKIP DGERVDFDDI HRKRMEKDLN

101 ELQALIEAHF ENRKKEEEEL VSLKDRIERR RAERAEQQRI RNEREKERQN

151 RL**A**EERARRE EEENRRKAED EARKKKALSN MMHFGGYIQK QAQTERKSGK

201 RQTEREKKKK IL**A**ERRKVLA IDHLNEDQLR EKAKELWQSI YNLEAEKFDL

251 QEKFKQQKYE INVLRNRIND NQKVSKTRGK AKVTGRWK

**Supplementary Table 2: Thin Filament Dynamics in 50mM HEPES, 25mM KCl, 4mM $MgCl_2$, 2mM $CaCl_2$**

| | Wildtype | | ΔE160 | |
|---|---|---|---|---|
| | **Lifetime** | **Transfer Efficiency** | **Lifetime** | **Transfer Efficiency** |
| **TnT** | 0.58 ± 0.13 | 0.71 ± 0.04 | 0.59 ± 0.15 | 0.74 ± 0.07 |
| **Troponin Complex** | 0.70 ± 0.12 | 0.59 ± 0.06 | 0.69 ± 0.10 | 0.59 ± 0.04 |
| **Thin Filament** | 0.68 ± 0.14 | 0.56 ± 0.07 | 0.71 ± 0.11 | 0.57 ±0.05 |
